# Ikzf1 association with Foxp3 for Foxp3-dependent gene repression in Treg cells: induction of autoimmunity and tumor immunity by disrupting the association

**DOI:** 10.1101/2023.08.12.553084

**Authors:** Kenji Ichiyama, Jia Long, Yusuke Kobayashi, Yuji Horita, Takeshi Kinoshita, Yamami Nakamura, Chizuko Kominami, Katia Georgopoulos, Shimon Sakaguchi

## Abstract

The transcription factor Foxp3 specifically expressed in regulatory T (Treg) cells controls Treg function by repressing some genes and activating others. We have shown here that the transcription factor Ikzf1 associates with Foxp3 via its exon 5 (called IkE5) and that conditional deletion of *IkE5* up-regulated the genes, including *Ifng*, normally repressed by Foxp3 upon TCR stimulation. *IkE5*-deletion in Treg cells indeed incurred IFN-γ overproduction, which destabilized Foxp3 expression and impaired suppressive function, consequently producing fatal systemic autoimmune diseases and evoking strong anti-tumor immunity. In addition, pomalidomide, which degrades IKZF1 and IKZF3, induced IFN-γ overproduction in human Treg cells. Mechanistically, the Foxp3/Ikzf1/Ikzf3 complex exerted gene-repressing function by competing with epigenetic co-activators, such as p300 and NFAT1, for binding to the target gene loci via chromatin remodeling. Collectively, the association of Ikzf1 with Foxp3 is essential for repressive function of Foxp3, and can be targeted to control autoimmunity and tumor immunity.

## INTRODUCTION

Regulatory T (Treg) cells are a functionally distinct CD4^+^ T-cell subset that plays crucial roles in maintaining immunological self-tolerance and homeostasis by suppressing aberrant or excessive immune responses.^1,2^ The transcription factor (TF) forkhead box protein P3 (Foxp3) is essential for the Treg cell function as illustrated by loss-of-function mutations of the *Foxp3* gene, which cause various immunological diseases such as autoimmune disease, allergy, and immunopathology in mice and humans through Treg cell deficiency or dysfunction.^3–8^ Foxp3 forms a large protein complex by interacting with a number of cofactors, including other TFs and epigenetic regulators.^9^ Upon T cell receptor (TCR) stimulation, the Foxp3 complex negatively controls the expression of some target genes, such as *Il2* and *Ifng,* as a strong repressor, while positively regulating other target genes, such as *Il2ra* and *Ctla4,* as an activator.^10–14^ Yet it is still contentious how a Foxp3 complex acts directly on its target genes as an activator, a repressor, or both, across diverse Treg cell states, and how the interactions of Foxp3 with other TFs, co-activators, and co-repressors affect Treg cell function and thereby cause autoimmune and other inflammatory diseases.^15–20^

The Ikaros TF family has five distinct members: Ikaros (encoded by *Ikzf1*), Helios (*Ikzf2*), Aiolos (*Ikzf3*), Eos (*Ikzf4*), and Pegasus (*Ikzf5*), all of which are characterized by two sets of highly conserved C2H2 zinc-finger motifs and critically involved in hematopoiesis and adaptive immunity.^21^ The first three or four motifs at the N-terminus are essential for regulating gene transcription through DNA binding, and the last two motifs at the C-terminus facilitate multimer formation as both homodimer and heterodimer with other family members.^22^ Ikzf family members, except Ikzf5, are highly expressed in Treg cells and physically associated with Foxp3.^9,20,23^ In particular, Helios and Eos have been reported to play a crucial role in the stability and suppressive function, respectively, of Treg cells.^23,24^ Interestingly, a recent attempt of comprehensive mutagenesis has suggested that variations of the Ikzf1-binding motifs impair Treg-specific chromatin accessibility.^25^ In addition, there is accumulating evidence that germline heterozygous mutations in *IKZF1*, especially in the exon 5 region of *IKZF1* (called IkE5), can cause immunodeficiency and autoimmune diseases in humans.^26–29^ These findings in mice and humans have prompted us to determine how Ikzf1 and Ikzf3 contribute to Treg cell function and how their anomalies, especially their interactions with Foxp3, are causative of autoimmune disease.

Here, we have shown that Ikzf1 is associated with Foxp3 via its IkE5 and that it forms a repressive Foxp3 complex together with Ikzf3, controlling genomic association of the complex to suppress the expression of the target genes including *Ifng*, which are normally repressed by Foxp3 upon TCR stimulation. Consequently, Treg-specific deletion of *IkE5* results in overproduction of IFN-γ in Treg cells, thereby impairs Foxp3 expression, hence Treg suppressive function, culminating in the development of fatal systemic autoimmune disease. The deletion can also evoke strong anti-tumor immunity. Our results show how a particular TF interacts with Foxp3 to exert repressive function and how the interaction can be pharmaceutically targeted to control immune responses.

## RESULTS

### Treg-specific deletion of *IkE5* causes fatal systemic autoimmunity

To identify the essential regions of Ikzf1 for association with Foxp3, we first generated several mutants that were deleted of C-terminal region (ΔC) or N-terminal region (ΔN) of Ikzf1, and conducted co-immunoprecipitation (Co-IP) assays (Figure 1A). Immunoblot analysis revealed that the ΔN mutant failed to interact with Foxp3 while the ΔC mutant did (Figure 1B). Since the deleted N-terminal region of Ikzf1 is composed of four exons, exon 4, exon 5 and exon 6/7, which contain four zinc-finger domains (Figure 1A), we next constructed mutants with deletion of each exon (ΔE4, ΔE5 and ΔE6/7) and examined which exon was required for the interaction with Foxp3 (Figure 1A). Co-IP assay showed that only ΔE5 mutant was unable to bind to Foxp3, indicating that IkE5 was essential for interaction with Foxp3 (Figure 1C).

**Figure 1.**
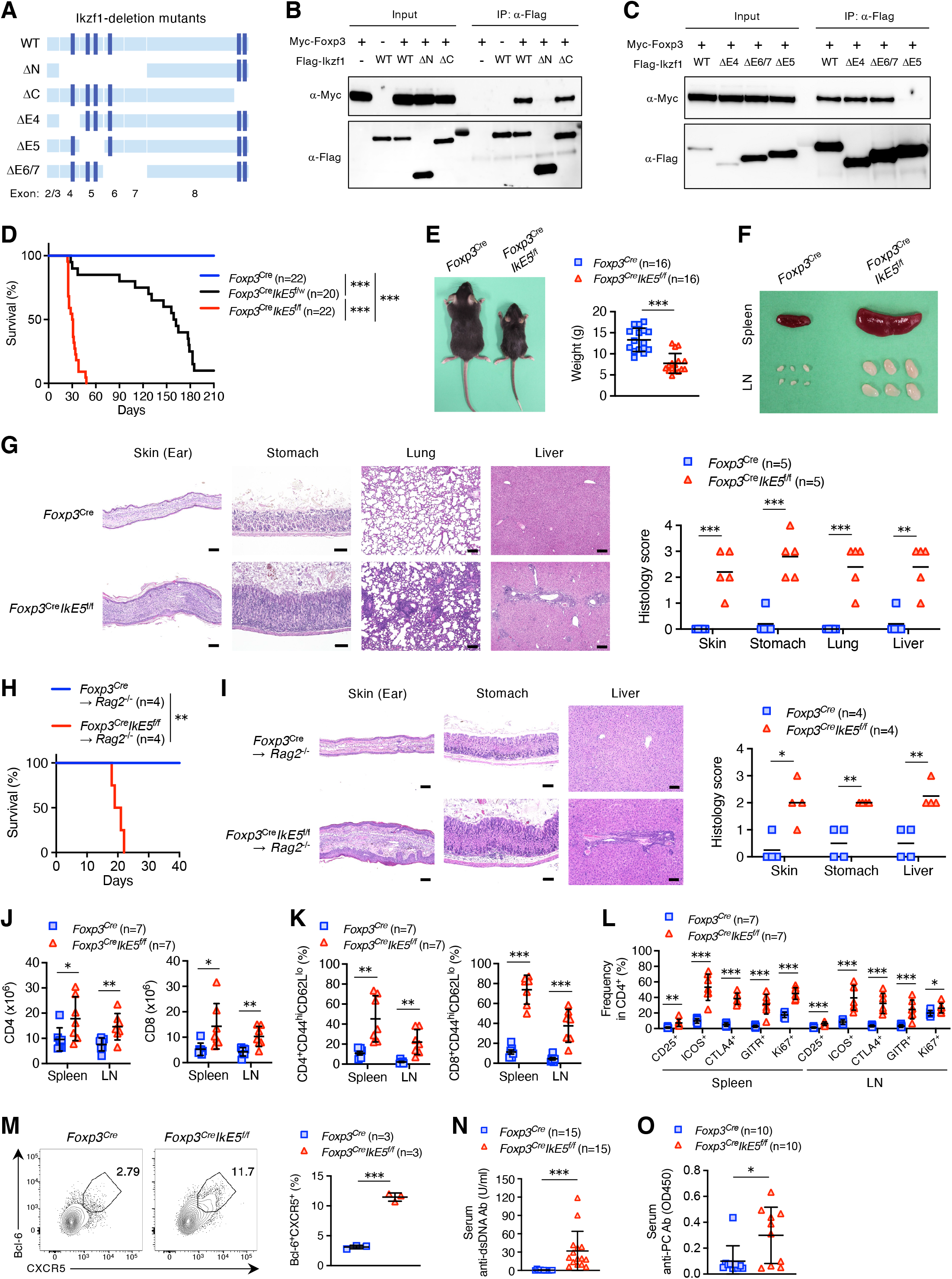
Treg-specific deletion of *IkE5* causes fatal systemic autoimmunity. (A) Schematic representation of the Ikzf1-deletion mutants constructed. Exons are shown as light blue boxes dark blue bars indicate zinc finger domains. (B) A representative image of the interaction between Foxp3 and Ikzf1 (WT) or Ikzf1-deletion mutants, which are deleted C-terminal region (ΔC) or N-terminal region (ΔN) of Ikzf1, in HEK293T cells by Co-IP. (C) A representative image of interaction between Foxp3 and Ikzf1 (WT) or Ikzf1-deletion mutants, which are deleted exon 4 (ΔE4), exon 5 (ΔE5) or exon 6/7 (ΔE6/7) of Ikzf1, in HEK293T cells by Co-IP. (D) Kaplan-Meier survival curve of *Foxp3*^Cre^ (*n* = 22), *Foxp3*^Cre^*IkE5*^f/w^ (*n* = 20) and *Foxp3*^Cre^*IkE5*^f/f^ mice (*n* = 22). (E) A representative appearance (left) and body weight (right) of *Foxp3*^Cre^ and *Foxp3*^Cre^*IkE5*^f/f^ mice at 3 to 4 weeks of age (mean ± SD, *n* = 16 per group). (F) A representative appearance of spleen and peripheral lymph nodes (LN) from *Foxp3*^Cre^ and *Foxp3*^Cre^*IkE5*^f/f^ mice at 3 to 4 weeks of age. (G) Hematoxylin and eosin (HE) staining of several tissues from *Foxp3*^Cre^ and *Foxp3*^Cre^*IkE5*^f/f^ mice at 3 to 4 weeks of age. Representative HE staining images (left) and histopathologic scoring (right) of indicated tissues from above mice are shown (*n* = 5 per group). Horizonal lines indicate mean. Scale bars, 100 μm. (H) Kaplan-Meier survival curve of CD45.1^+^*Rag2*^−/−^ mice transferred with splenocytes (3 × 10^7^) from either *Foxp3*^Cre^ or *Foxp3*^Cre^*IkE5*^f/f^ mice (*n* = 4 per group). (I) HE staining of several tissues from CD45.1^+^*Rag2*^−/−^ mice transferred as indicated in (H). Representative HE staining images (left) and histopathologic scoring (right) of indicated tissues from above mice are shown (*n* = 4 per group). Horizonal lines indicate mean. Scale bars, 100 μm. (J) Absolute numbers of CD4^+^ and CD8^+^ T cells in the spleen and LN from *Foxp3*^Cre^ and *Foxp3*^Cre^*IkE5*^f/f^ mice at 3 to 4 weeks of age (mean ± SD, *n* = 7 per group). (K) Frequencies of CD4^+^CD44^hi^CD62L^lo^ and CD8^+^CD44^hi^CD62L^lo^ T cells in the spleen and LN from *Foxp3*^Cre^ and *Foxp3*^Cre^*IkE5*^f/f^ mice at 3 to 4 weeks of age (mean ± SD, *n* = 7 per group). (L) Frequencies of indicated molecules-expressing CD4^+^ T cells in the spleen and LN from *Foxp3*^Cre^ and *Foxp3*^Cre^*IkE5*^f/f^ mice at 3 to 4 weeks of age (mean ± SD, *n* = 7 per group). (M) Frequencies of CD4^+^Bcl-6^+^CXCR5^+^ T (Tfh) cells in the spleen from *Foxp3*^Cre^ and *Foxp3*^Cre^*IkE5*^f/f^ mice at 3 to 4 weeks of age (right)(mean ± SD, *n* = 3 per group). A representative flow cytometry plot of CD4^+^ splenocytes in the above mice (left). (N) Concentration of anti-dsDNA antibody in serum of *Foxp3*^Cre^ and *Foxp3*^Cre^*IkE5*^f/f^ mice at 3 to 4 weeks of age, determined by ELISA (mean ± SD, *n* = 15 per group). (O) Concentration of anti-PC antibody in serum of *Foxp3*^Cre^ and *Foxp3*^Cre^*IkE5*^f/f^ mice at 3 to 4 weeks of age, determined by ELISA (mean ± SD, *n* = 10 per group). Data are representative of at least three independent experiments (B,C,F) or summary of at least two independent experiments (D,E,G-O). *P* values determined by two-tailed unpaired *t*-tests (E,M-O), log-rank test (D,H) or unpaired *t*-tests followed by Holm-Šídák multiple comparisons test (G,I-L). \**P*< 0.05; \*\**P* < 0.01; \*\*\**P* < 0.001. Also see Figure S1.

To investigate the role of interaction between Ikzf1 and Foxp3 in Treg cell biology *in vivo*, we generated *Foxp3*^Cre^*IkE5*^f/f^ mice by crossing mice in which *IkE5* was flanked with loxP to *Foxp3*^IRES-YFP-Cre^ mice expressing yellow fluorescent protein (YFP)-Cre recombinase fusion protein under the control of Foxp3 regulatory elements.^30,31^ In these *Foxp3*^Cre^*IkE5*^f/f^ mice, we observed efficient Treg-specific deletion of *IkE5* (Figure S1A). Furthermore, Co-IP analysis confirmed the dissociation of Ikzf1 and Foxp3 interaction in *IkE5*-deficient Treg cells (Figure S1B). *Foxp3*^Cre^*IkE5*^f/f^ mice died between 20 and 50 days after birth (Figure 1D). Of note, *Foxp3*^Cre^*IkE5*^f/w^ mice developed less severe but still lethal autoimmunity as seen in haploinsufficiency of human germline *IKZF1* mutations (Figure 1D).^26–29^ *Foxp3*^Cre^*IkE5*^f/f^ mice showed reduced body weight, lymphadenopathy and splenomegaly, and massive infiltrations of leukocytes into skin (ear), stomach, lung and liver (Figures 1E-1G). Moreover, fatal inflammation and tissue lesions could be adoptively transferred by splenocytes from *Foxp3*^Cre^*IkE5*^f/f^ mice into syngeneic *Rag2*^−/−^ mice, indicating autoimmune nature of the inflammation (Figures 1H and 1I).

The frequencies and absolute numbers of both CD4^+^ and CD8^+^ T cells, in particular, CD44^hi^CD62L^lo^ effector memory cells, were markedly increased in peripheral lymphoid organs of *Foxp3*^Cre^*IkE5*^f/f^ mice (Figures 1J and 1K; Figure S1C). CD4^+^Foxp3^−^ T cells in *Foxp3*^Cre^*IkE5*^f/f^ mice exhibited enhanced expressions of activation markers, including CD25, inducible T-cell co-stimulator (ICOS), cytotoxic T-lymphocyte associated protein 4 (CTLA4) and glucocorticoid-induced TNFR-related protein (GITR) as well as the proliferation marker Ki67 (Figure 1L). Similarly, T cells producing cytokines, such as interferon-γ (IFN-γ) and interleukin-4 (IL-4), increased in *Foxp3*^Cre^*IkE5*^f/f^ mice (Figure S1D).

The frequencies of both CD4^+^Bcl-6^+^CXCR5^+^ follicular helper T (Tfh) cells and IgM^+^GL7^+^ germinal center B (GCB) cells were increased in *Foxp3*^Cre^*IkE5*^f/f^ mice, suggesting their contribution to autoantibody production (Figures 1M and S1E). Consistently, the serum concentrations of IgG, IgM and IgE were all markedly increased in *Foxp3*^Cre^*IkE5*^f/f^ mice (Figure S1F). Of note, *Foxp3*^Cre^*IkE5*^f/f^ mice showed enhanced production of serum autoantibodies against double-stranded DNA (dsDNA) and gastric parietal cells (PC), which are indicative of systemic and organ-specific autoimmunity, respectively (Figures 1N and 1O). Collectively, Treg-specific deletion of *IkE5* produced fatal systemic autoimmune and inflammatory diseases, which were similar to scurfy disease due to a loss-of-function mutation in the *Foxp3* gene.^32^

### *IkE5*-deficient Treg cells show impaired suppressive function and functional instability

With the autoimmunity in *Foxp3*^Cre^*IkE5*^f/f^ mice, we assessed the role of *IkE5* in Treg cell function and maintenance. To exclude possible indirect effects by inflammation, we made use of *Foxp3*^Cre/+^*IkE5*^f/f^ mosaic female mice, which possessed both YFP^+^ *IkE5*-deficient Treg cells and YFP^−^ *IkE5*-intact Treg cells as a result of random inactivation of X chromosome-linked genes. *Foxp3*^Cre/+^*IkE5*^f/f^ mice indeed showed no overt signs of autoimmunity or body weight loss during 4 weeks of observation, with no alteration in the expression of activation markers, such as CD25, ICOS, CTLA4, GITR and Ki67, in YFP^+^ Treg cells (Figures S2A-S2C). However, the ratio between YFP^+^Foxp3^+^ and YFP^−^Foxp3^+^ Treg populations in *Foxp3*^Cre/+^*IkE5*^f/f^ mice was significantly decreased compared to the corresponding ratio in *Foxp3*^Cre/+^ mice (Figure 2A). There was no significant changes in the ratio in the thymus (Figure S2D). The results suggested that *IkE5*-deletion led to a decline in Treg cell survival or their Foxp3 expression. To examine then whether *IkE5*-deficient Treg cells would lose Foxp3 expression, we isolated YFP^+^ Treg cells from *Foxp3*^Cre/+^ and *Foxp3*^Cre/+^*IkE5*^f/f^ mice and transferred each of them into lymphopenic *Rag2*^−/−^ mice. Compared with wild-type Treg cells, transfer of *IkE5-*deficient Treg cells resulted in a significant increase of Foxp3^−^ (exTreg) donor cells in the peripheral lymphoid organs (Figure 2B). To further confirm the instability of Treg cells by *IkE5*-deficiency in the steady state, we generated Foxp3 fate-mapping mice by crossing *Foxp3*^Cre/+^*IkE5*^f/f^ mice to *Rosa26*^rfp^ reporter mice inserted at the *ROSA26* locus with a loxP site-flanked STOP cassette before a DNA sequence encoding red fluorescent protein (RFP).^33^ The frequency of YFP^−^RFP^+^ exTreg cells in the peripheral lymphoid organs of *Foxp3*^Cre/+^*IkE5*^f/f^*R26*^rfp/+^ mice were indeed increased significantly compared to *Foxp3*^Cre/+^*R26*^rfp/+^ mice (Figure 2C). Regarding this Treg instability, Kim et al previously reported that *Ikzf2*-deficient Treg cells were unstable in Foxp3 expression because of a diminished activation of the STAT5 pathway.^24^ It is also known that the DNA hypomethylation pattern of Treg-specific demethylation regions (TSDRs) is required for the maintenance of Foxp3 expression.^13,34^ However, DNA demethylation of the *Foxp3* CNS2 region as well as the expression of phosphorylated STAT5 (pSTAT5) were normal in *IkE5*-deficient Treg cells (Figures S2E and S2F).

**Figure 2.**
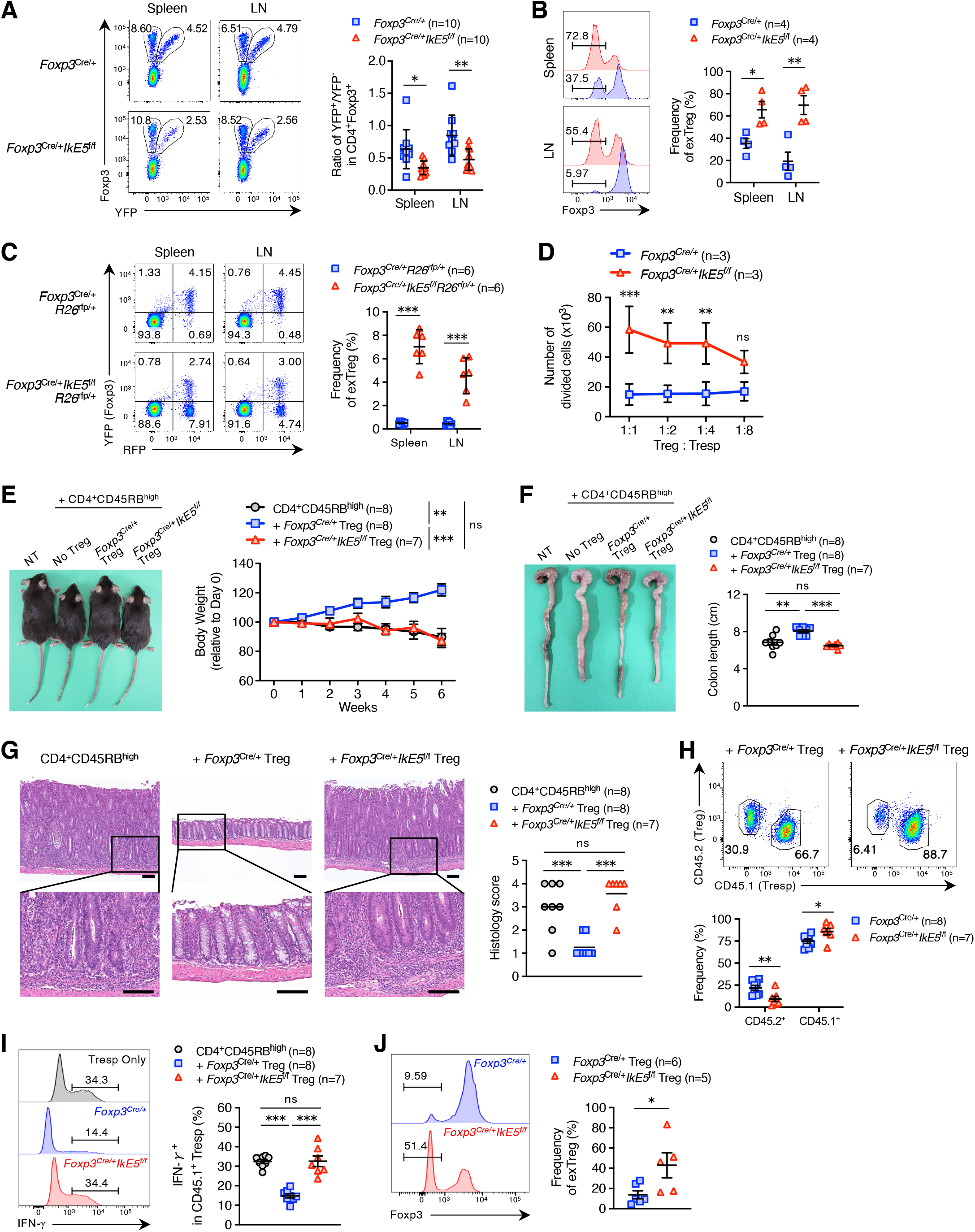
*IkE5*-deficient Treg cells show impaired suppressive function and functional instability. (A) A representative flow cytometry plot of CD4^+^ T cells (left) and the ratio of YFP^+^/YFP^−^ Treg cells (right) in the spleen and LN from *Foxp3*^Cre/+^ and *Foxp3*^Cre/+^*IkE5*^f/f^ mice at 3 to 4 weeks of age (mean ± SD, *n* = 10 per group). (B) A representative flow cytometry histogram of Foxp3 in CD4^+^CD45.2^+^ cells (left) and frequencies of CD4^+^CD45.2^+^Foxp3^−^ (exTreg) cells (right) in the spleen and LN from CD45.1^+^*Rag2*^−/−^ mice transferred with CD4^+^CD45.2^+^YFP^+^ Treg cells (1 × 10^5^) from either *Foxp3*^Cre/+^ (*n* = 4) or *Foxp3*^Cre/+^*IkE5*^f/f^ (*n* = 4) mice at day 8 after the transfer (mean ± SEM). (C) A representative flow cytometry plot of CD4^+^ T cells (left) and frequencies of CD4^+^YFP^−^RFP^+^ T (exTreg) cells (right) in the spleen and LN from *Foxp3*^Cre/+^*R26*^rfp/+^ and *Foxp3*^Cre/+^*IkE5*^f/f^*R26*^rfp/+^ mice at 3 to 4 weeks of age (mean ± SD, *n* = 6 per group). (D) CTV-labeled Purified CD4^+^CD45.1^+^ naïve T (Tresp) cells (5 × 10^4^) were cultured with CD3^+^ T cell-depleted splenocytes (1 × 10^5^) and CD4^+^CD45.2^+^FYP^+^ Treg cells from *Foxp3*^Cre/+^ or *Foxp3*^Cre/+^*IkE5*^f/f^ mice at the indicated Treg : Tresp ratios in the presence of anti-CD3 antibody (1 μg/ml) for 3 days. Absolute numbers of Tresp cells were assayed with flow cytometry (mean ± SD, *n* = 3 per group). (E) A representative appearance of non-transferred *Rag2*^−/−^ mice (NT) and recipient *Rag2*^−/−^ mice transferred with CD4^+^CD45.1^+^CD45RB^hi^ naïve T cells (1 × 10^5^) alone (*n* = 8)(black) or along with CD4^+^CD45.2^+^YFP^+^ Treg cells (1 × 10^5^) from either *Foxp3*^Cre/+^ (*n* = 8)(blue) or *Foxp3*^Cre/+^*IkE5*^f/f^ mice (*n* = 7)(red) at 6 weeks after transfer (left). Kinetics of body weight changes of *Rag2*^−/−^ recipients over 6 weeks after transfer (right)(mean ± SEM). (F) A representative appearance (left) and length (right) of colons from non-transferred *Rag2*^−/−^ mice (NT) and *Rag2*^−/−^ recipients transferred as indicated in (E) at 6 weeks after transfer (mean ± SEM). (G) A representative HE staining image of colons from *Rag2*^−/−^ recipients transferred as indicated in (E) at 6 weeks after transfer (left). Magnification is x100 (top) and x300 (bottom). Histopathologic scoring of colons from above recipients (right). Horizontal lines indicate mean. (H) A representative flow cytometry plot of CD4^+^ T cells (up) and frequencies of CD4^+^CD45.2^+^ Treg and CD4^+^CD45.1^+^ Tresp cells (bottom) in LN from *Rag2*^−/−^ recipients transferred as indicated in (E) at 6 weeks after transfer (mean ± SEM). (I) Representative flow cytometry histograms of IFN-γ in CD4^+^CD45.1^+^ Tresp cells (left) and frequencies of CD4^+^CD45.1^+^IFN-γ ^+^ Tresp cells (right) in LN from *Rag2*^−/−^ recipients transferred as indicated in (E) at 6 weeks after transfer (mean ± SEM). (J) Representative flow cytometry histograms of Foxp3 (left) and frequency of exTreg cells (right) in CD4^+^CD45.2^+^ Treg cells in LN from *Rag2*^−/−^ recipients transferred as indicated in (E) at 6 weeks after transfer (mean ± SEM). Data are summary of at least three independent experiments (A-J). *P* values determined by unpaired *t*-tests followed by Holm-Šídák multiple comparisons test (A-C), two-way ANOVA followed by Šídák multiple comparisons test (D), two-way ANOVA followed by Tukey’s multiple comparisons test (E), ordinary one-way ANOVA followed by Tukey’s multiple comparisons test (F,G,I) or two-tailed unpaired *t*-tests (H,J). ns, not significant, \**P*< 0.05; \*\**P* < 0.01; \*\*\**P* < 0.001. Also see Figure S2.

We next assessed the role of *IkE5* in suppressive function of Treg cells. In an *in vitro* suppression assay, *IkE5*-deficient Treg cells failed to suppress the numerical expansion of responder T cells compared to their wild-type counterparts (Figure 2D). To assess *in vivo* suppressive activity, we used the T-cell transfer model of colitis: Treg cells from CD45.2^+^*Foxp3*^Cre/+^ or CD45.2^+^*Foxp3*^Cre/+^*IkE5*^f/f^ mice were transferred into *Rag2*^−/−^ mice together with congenic CD45.1^+^CD4^+^CD25^−^CD45RB^hi^ naïve T cells.^35^ As shown in Figures 2E-2G, while wild-type Treg cells prevented body weight loss, change in colon length, and colonic inflammation, *IkE5*-deficient Treg cells failed to exhibit such suppressive effects. Further, *Rag2*^−/−^ mice recipient of *IkE5*-deficient Treg cells showed a significant increase of responder CD4^+^ T cells, especially those producing IFN-γ (Figures 2H and 2I). Notably, a significant portion of the transferred *IkE5*-deficient Treg cells was converted to exTreg cells when compared to similarly transferred wild-type Treg cells (Figure 2J), suggesting severely compromised functional stability in *IkE5*-deficient Treg cells. These results collectively indicate that the IkE5 region is critically required for the suppressive function and functional stability of Treg cells.

### Treg-specific deletion of *IkE5* evokes strong anti-tumor immunity

We next investigated whether the *IkE5*-deletion in Treg cells would facilitate not only autoimmunity but also anti-tumor immunity. By crossing *IkE5*^f/f^ mice to *Foxp3*^eGFP-Cre-ERT2^ mice,^36^ we established *Foxp3*^eGFP-Cre-ERT2^*IkE5*^f/f^ mice that allowed tamoxifen-inducible Treg-specific deletion of *IkE5*. We inoculated into the mice B16F0 murine melanoma or MC38 murine colon adenocarcinoma cells intradermally, and injected tamoxifen intraperitoneally on day 0, 1 and 3 (Figure 3A). The growth of both B16F0 and MC38 cells was strongly inhibited in *Foxp3*^eGFP-Cre-ERT2^*IkE5*^f/f^ mice compared to control *Foxp3*^eGFP-Cre-ERT2^ mice (Figures 3B and 3C; Figures S3A and S3B). *Foxp3*^eGFP-Cre-ERT2^*IkE5*^f/f^ mice showed a profound reduction of Treg cells in the draining lymph nodes (dLNs) and tumor tissues, but not in non-draining lymph nodes (ndLNs) when examined on day 10 after tumor cell inoculation (Figure 3D). Moreover, the ratio of intratumoral CD8^+^ T to Treg cells was markedly increased in *Foxp3*^eGFP-Cre-ERT2^*IkE5*^f/f^ mice (Figure 3E), implying strong anti-tumor responses. Consistently, a higher frequency of intratumoral CD4^+^Foxp3^−^ T cells in *Foxp3*^eGFP-Cre-ERT2^*IkE5*^f/f^ mice produced anti-tumor effector cytokines, including IFN-γ and tumor necrosis factor-α (TNF-α) (Figure 3F). Taken together, IkE5 is required for the suppressive function and functional stability of Treg cells in the tumor microenvironment.

**Figure 3.**
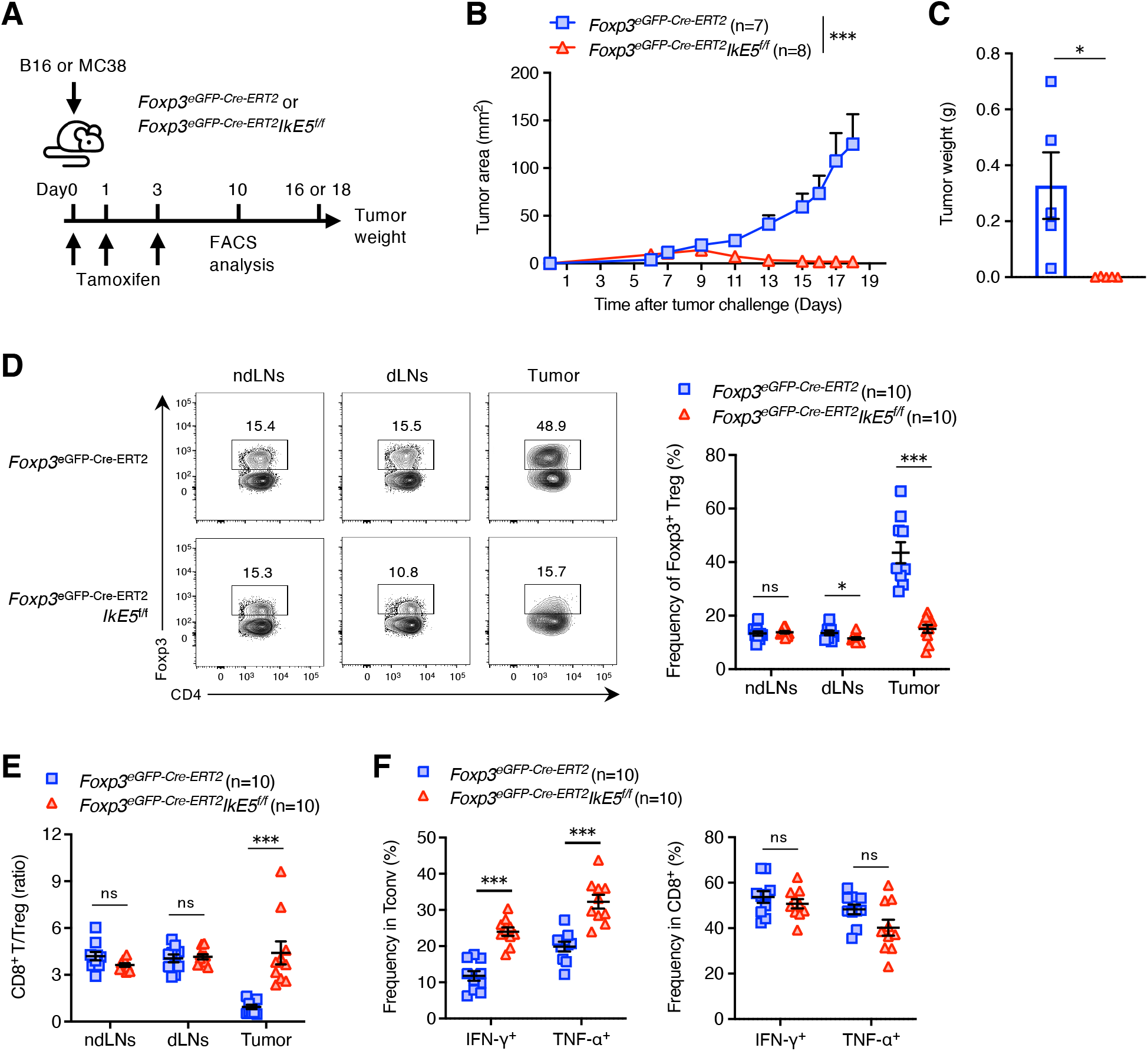
Treg-specific deletion of *IkE5* evokes strong anti-tumor immunity. (A) Schematic representation of procedure for murine tumor models conducted. (B) *Foxp3*^eGFP-Cre-ERT2^ (*n* = 7) and *Foxp3*^eGFP-Cre-ERT2^*IkE5*^f/f^ (*n* = 8) mice were intradermally inoculated with MC38 murine colon adenocarcinoma cells (2 × 10^5^) into their shaved flanks (Day 0) and intraperitoneally injected with tamoxifen on day 0, 1 and 3. Tumor area (mm^2^) was measured over 18 days (mean ± SEM). (C) Tumor weight in *Foxp3*^eGFP-Cre-ERT2^ and *Foxp3*^eGFP-Cre-ERT2^*IkE5*^f/f^ mice inoculated as indicated in (B) was measured on 18 days (mean ± SEM, *n* = 5 per group). (D) Representative flow cytometry plots of CD4^+^ T cells (left) and frequencies of CD4^+^Foxp3^+^ Treg cells (right) in Tumor, dLNs and ndLNs from each mouse inoculated as indicated in (B) at 10 days (mean ± SEM, *n* = 10 per group). (E) Ratio of CD8^+^/CD4^+^Foxp3^+^ Treg cells in Tumor, dLNs and ndLNs from each mice inoculated as indicated in (B) at 10 days (mean ± SEM, *n* = 10 per group). (F) Frequencies of indicated cytokines-producing Tconv or CD8^+^ T cells in Tumor from each mouse inoculated as indicated in (B) at 10 days (mean ± SEM, *n* = 10 per group). Data are summary of at least two independent experiments (B-F). *P* values determined by ordinary two-way ANOVA followed by Šídák multiple comparisons test (B) or two-tailed unpaired *t*-tests (C) or unpaired *t*-tests followed by Holm-Šídák multiple comparisons test (D-F). ns, not significant, \**P*< 0.05; \*\**P*< 0.01; \*\*\**P* < 0.001. Also see Figure S3.

### IkE5 is required for Foxp3-dependent gene repression in Treg cells

To explore the molecular program affected by *IkE5*-deletion in Treg cells, we performed RNA-sequencing (RNA-seq) analysis of FYP^+^ Treg cells from *Foxp3*^Cre/+^ and *Foxp3*^Cre/+^*IkE5*^f/f^ mice. Principal component analysis (PCA) clearly separated wild-type and *IkE5*-deficient Treg cells based on both location and genotype (Figure S4A), indicating a significant alteration in the transcriptome pattern in *IkE5*-deleted Treg cells. Differential gene expression analysis showed that 469 genes were up-regulated (Up) and 248 genes were down-regulated (Down) in *IkE5*-deficient Treg cells compared to wild-type Treg cells (Figure 4A and Table S1). Since Up genes contained a number of Treg-signature genes (Figure 4A), we next performed a gene set enrichment analysis (GSEA) and a cumulative distribution function (CDF) analysis on the RNA-seq data. Both analyses revealed that the “Treg down genes”^37^ were more highly expressed in *IkE5*-deficient Treg cells compared to wild-type Treg cells, while the “Treg up genes”^37^ were comparable between them (Figures 4B and 4C). Moreover, when “Foxp3 dependent genes”^38^ were compared, they were mainly up-regulated in *IkE5*-deficient Treg cells (Figures 4D and 4E), suggesting impaired Foxp3-dependent gene repression in *IkE5*-deleted Treg cells. In agreement with these findings, we confirmed that the expression of Treg down-regulated genes, such as *Il2*, *Ifng*, *Il4*, *Il21*, *Tbx21*, *Cd4*, *Plscr1* and *Zap70*, were significantly increased in *IkE5*-deficient Treg cells (Figure 4F). We further confirmed at the protein level the enhancement of IL-2 and IFN-γ production from *IkE5*-deficient Treg cells (Figure 4G). In contrast, the expression of Treg up-regulated genes, such as *Ctla4*, *Ikzf2*, *Ikzf4*, *Itgae*, *Nt5e* and *Il10*, were mostly normal, while the expression of some genes, e.g., *Il2ra* and *Tnfrsf18*, were further enhanced in *IkE5*-deficient Treg cells at both the mRNA and protein levels (Figures 4F and S4B). In addition to Treg-signature genes, GSEA revealed an enhanced expression of gene signatures related to “Inflammatory response”, “IL-2 STAT5 signaling” and “IFN-gamma response” in *IkE5*-deficient Treg cells (Figures 4H and 4I). Collectively, these results suggest that IkE5 plays a critical role in Foxp3-dependent gene repression to maintain Treg cell homeostasis.

**Figure 4.**
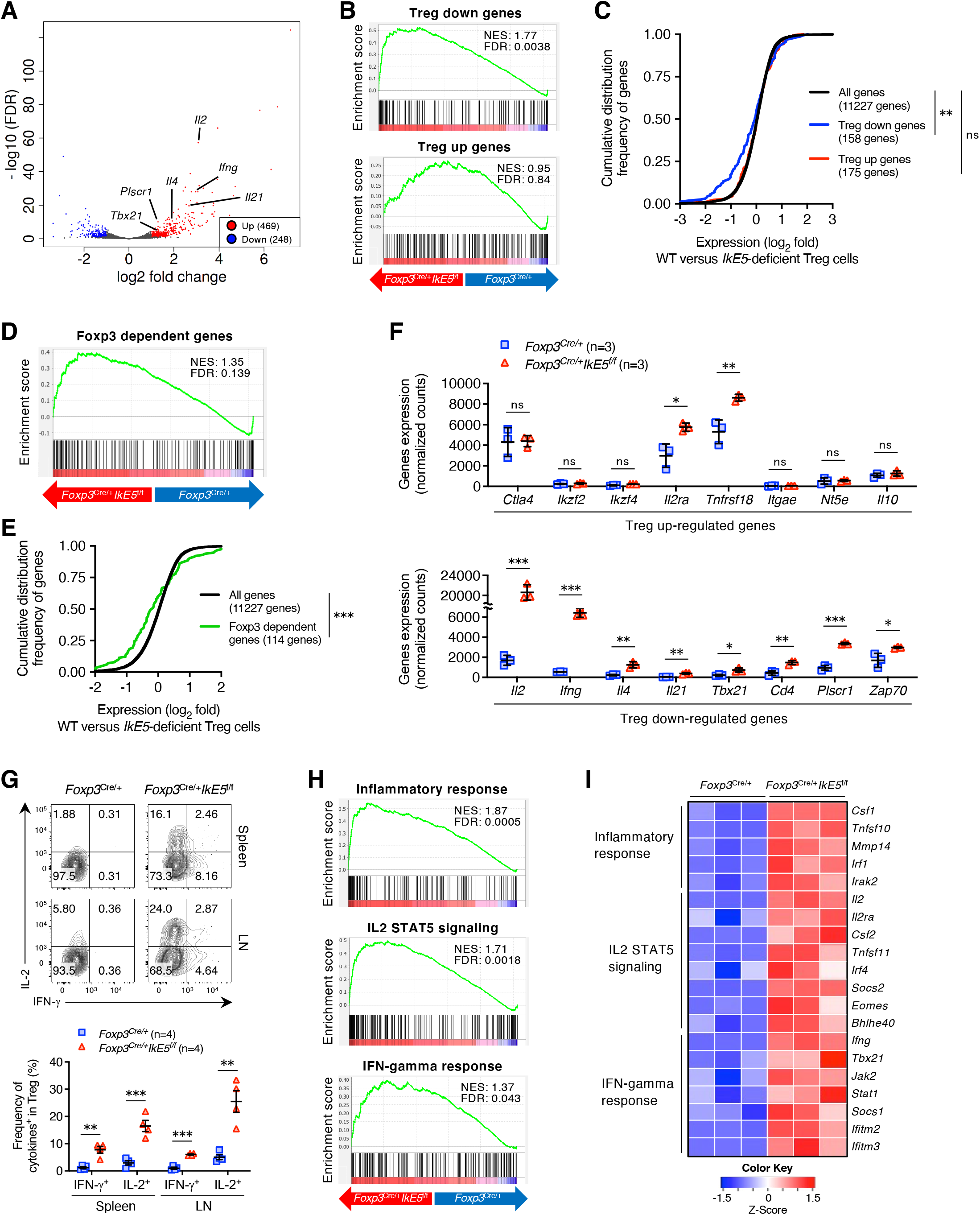
IkE5 is required for Foxp3-dependent gene repression in Treg cells. (A) RNA-seq analysis of CD4^+^YFP^+^ Treg cells from *Foxp3*^Cre/+^ or *Foxp3*^Cre/+^*IkE5*^f/f^ mice at 3 to 4 weeks of age. Volcano-plot comparing the expression of genes in wild-type and *IkE5*-deficient Treg cells. Transcripts Per Kilobase Million (TPM) were averaged from three biological replicates. Differential Expressed Genes (DEGs)(fold change, 2-fold; FDR < 0.1) are shown in red (Up) and blue (Down). Exact numbers of DEGs and name of representative genes are described. (B) Ranked enrichment plots of “Treg down genes” (up) and “Treg up genes” (bottom) in gene set enrichment analysis (GSEA) of *Foxp3*^Cre/+^*IkE5*^f/f^ versus *Foxp3*^Cre/+^ Treg cells. (C) Cumulative distribution function (CDF) analysis of the gene expression in wild-type and *IkE5*-deficient Treg cells for “Treg down genes” and “Treg up genes”. (D) Ranked enrichment plot of “Foxp3 dependent genes” in GSEA of *Foxp3*^Cre/+^*IkE5*^f/f^ versus *Foxp3*^Cre/+^ Treg cells. (E) CDF analysis of the gene expression in wild-type and *IkE5*-deficient Treg cells For “Foxp3 dependent genes”. (F) Normalized read counts of selected Treg up-regulated genes (up) and Treg down-regulated genes (bottom) in CD4^+^YFP^+^ Treg cells from *Foxp3*^Cre/+^ or *Foxp3*^Cre/+^*IkE5*^f/f^ mice (mean ± SD, *n* = 3 per group). (G) A representative flow cytometry plot of CD4^+^YFP^+^ Treg cells (up) and frequencies of indicated cytokine-producing Treg cells (bottom) in the spleen and LN from *Foxp3*^Cre/+^ and *Foxp3*^Cre/+^*IkE5*^f/f^ mice at 3 to 4 weeks of age (mean ± SEM, *n* = 4 per group). (H) Ranked enrichment plots of “Inflammatory response”, “IL-2 STAT5 signaling” and “IFN-gamma response” in GSEA of *Foxp3*^Cre/+^*IkE5*^f/f^ versus *Foxp3*^Cre/+^ Treg cells. (I) Heatmap of selective genes related to “Inflammatory response”, “IL-2 STAT5 signaling” and “IFN-gamma response” in GSEA (*n* = 3 per group). Data are representative of two independent experiments (A-F,H,I) or summary of three independent experiments (G). *P* values determined by Kolmogorov-Smirnov test (C,E) or unpaired *t*-tests followed by Holm-Šídák multiple comparisons test (F,G). ns, not significant, \**P*< 0.05; \*\**P*< 0.01; \*\*\**P* < 0.001. Also see Figure S4.

### Overproduction of IFN-γ promotes the instability and dysfunction of *IkE5*-deficient Treg cells

Since *IkE5*-deficient Treg cells highly produced IL-2 and IFN-γ, we next generated mice deficient of either cytokine and examined whether overproduction of these cytokines was causative of fatal inflammation in *Foxp3*^Cre^*IkE5*^f/f^ mice. *Foxp3*^Cre^*IkE5*^f/f^*Ifng*^−/−^ mice showed significant prolongation of survival compared to *Foxp3*^Cre^*IkE5*^f/f^ mice, in contrast with no alteration in the survival of *Foxp3*^Cre^*IkE5*^f/f^*Il2*^f/f^ mice (Figures 5A and S5). Since IFN-γ is known to drive Treg cells to be fragile and reduce their suppressive activity,^39^ we generated *Foxp3*^Cre/+^*IkE5*^f/f^*Ifng*^−/−^*R26*^rfp/+^ mice and examined whether IFN-γ deficiency would repair the instability of *IkE5*-deficient Treg cells. The IFN-γ deficiency indeed significantly inhibited the increase of YFP^−^ RFP^+^ exTreg cells in the peripheral lymphoid organs of *Foxp3*^Cre/+^*IkE5*^f/f^*R26*^rfp/+^ mice (Figure 5B). Moreover, transfer of *IkE5* and *Ifng* double*-*deficient Treg cells into lymphopenic *Rag2*^−/−^ mice showed their much less conversion to exTreg cells compared with similar transfer of *IkE5*-deficient Treg cells (Figure 5C). To further substantiate the role of IFN-γ for the instability of *IkE5*-deficient Treg cells, we cultured FYP^+^ Treg cells from *Foxp3*^Cre/+^ or *Foxp3*^Cre/+^*IkE5*^f/f^ mice under TCR and IL-2 stimulation with/without neutralizing anti-IFN-γ antibody for 7 days *in vitro*. While about 35% of *IkE5*-deficient Treg cells lost Foxp3 expression, anti-IFN-γ antibody prevented the Foxp3 loss (Figure 5D).

**Figure 5.**
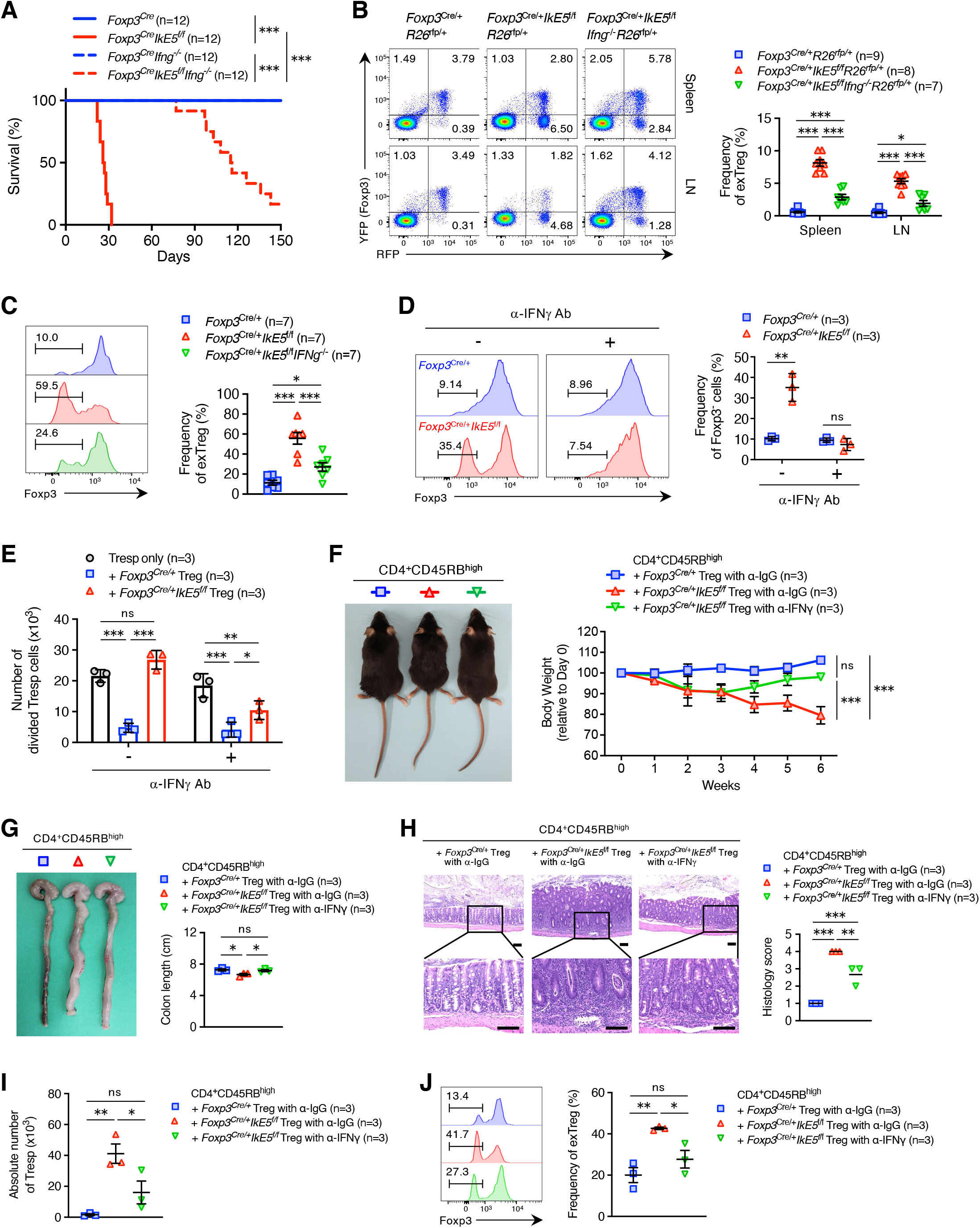
Overproduction of IFN-γ promotes the instability and dysfunction of *IkE5*-deficient Treg cells. (A) Kaplan-Meier survival curve of *Foxp3*^Cre^, *Foxp3*^Cre^*IkE5*^f/f^, *Foxp3*^Cre^*Ifng*^−/−^ and *Foxp3*^Cre^*IkE5*^f/f^*Ifng*^−/−^ mice (*n* = 12 per group). (B) A representative flow cytometry plot of CD4^+^ T cells (left) and frequencies of CD4^+^YFP^−^RFP^+^ Treg (exTreg) cells (right) in the spleen and LN from *Foxp3*^Cre/+^*R26*^rfp/+^ (*n* = 9), *Foxp3*^Cre/+^*IkE5*^f/f^*R26*^rfp/+^ (*n* = 8) and *Foxp3*^Cre/+^*IkE5*^f/f^*Ifng*^−/−^*R26*^rfp/+^ (*n* = 7) mice at 3 to 4 weeks of age (mean ± SEM). (C) A representative flow cytometry histogram of Foxp3 in CD4^+^CD45.2^+^ cells (left) and frequencies of CD4^+^CD45.2^+^Foxp3^−^ (exTreg) cells (right) in the spleen and LN from CD45.1^+^*Rag2*^−/−^ mice transferred with CD4^+^CD45.2^+^YFP^+^ Treg cells (1 × 10^5^) from either *Foxp3*^Cre/+^ (*n* = 4) or *Foxp3*^Cre/+^*IkE5*^f/f^ (*n* = 4) mice at day 7 after the transfer (mean ± SEM). (D) Purified CD4^+^FYP^+^ Treg cells (1 × 10^5^) from *Foxp3*^Cre/+^ and *Foxp3*^Cre/+^*IkE5*^f/f^ were cultured with Dynabeads mouse CD3/CD28 T cell stimulator (25 μl/ml) and IL-2 (100 U/ml) in the presence of anti-IFN-γ or anti-IgG antibody (10 μg/ml) for 7 days. A representative flow cytometry histogram of Foxp3 (left) and frequencies of Foxp3^−^ population (right) in cultured CD4^+^ T cells (mean ± SD, *n* = 3 per group). (E) CTV-labeled Purified CD4^+^CD45.1^+^ naïve T (Tresp) cells (5 × 10^4^) were cultured with CD3^+^ T cell-depleted splenocytes (1 × 10^5^) and CD4^+^CD45.2^+^FYP^+^ Treg cells from *Foxp3*^Cre/+^ or *Foxp3*^Cre/+^*IkE5*^f/f^ mice at 1 : 2 Treg/Tresp ratio in the presence of anti-CD3 antibody (1 μg/ml) and anti-IFN-γ (10 μg/ml) or anti-IgG antibody (10 μg/ml) for 3 days. Absolute numbers of Tresp cells were assayed with flow cytometry (mean ± SD, *n* = 3 per group). (F) CD4^+^CD45.1^+^CD25^−^CD45RB^hi^ naïve T cells (1 × 10^5^) and CD4^+^CD45.2^+^FYP^+^ Treg cells (1 × 10^5^) from *Foxp3*^Cre/+^ or *Foxp3*^Cre/+^*IkE5*^f/f^ mice were mixed and intravenously transferred into *Rag2*^−/−^ recipients. For IFN-γ neutralization, anti-IFN-γ antibody (250 μg/mouse) intraperitoneally injected into *Rag2*^−/−^ recipients transferred with *IkE5*-deficient Treg cells (green) every 3 days. As control, anti-IgG antibody (250 μg/mouse) intraperitoneally injected into *Rag2*^−/−^ recipients transferred with wild-type (blue) or *IkE5*-deficient Treg cells (red) every 3 days. A representative appearance of recipient *Rag2*^−/−^ mice transferred as indicated above at 6 weeks after transfer (left). Kinetics of body weight changes of recipient *Rag2*^−/−^ mice over 6 weeks after transfer (right)(mean ± SEM, *n* = 3 per group). (G) A representative appearance (left) and length (right) of colons from *Rag2*^−/−^ recipients transferred as indicated in (F) at 6 weeks after transfer (mean ± SEM, *n* = 3 per group). (H) Representative HE staining images of colons from *Rag2*^−/−^ recipients transferred as indicated in (F) at 6 weeks after transfer (left). Magnification is x100 (top) and x300 (bottom). Histopathologic scoring of colons from above recipients (right). Horizontal lines indicate mean. (I) Absolute number of CD4^+^CD45.1^+^ Tresp cells in LN from *Rag2*^−/−^ recipients transferred as indicated in (F) at 6 weeks after transfer (mean ± SEM, *n* = 3 per group). (J) A representative flow cytometry histogram of Foxp3 (left) and frequency of exTreg cells (right) in CD4^+^CD45.2^+^ Treg cells in LN from *Rag2*^−/−^ recipients transferred as indicated in (F) at 6 weeks after transfer (mean ± SEM, *n* = 3 per group). Data are summary of at least three independent experiments (A-E) or representative of two independent experiments (F-J). *P* values determined by log-rank test (A), unpaired *t*-tests followed by Holm-Šídák multiple comparisons test (D), two-way ANOVA followed by Tukey’s multiple comparisons test (B,E,F) or ordinary one-way ANOVA followed by Tukey’s multiple comparisons test (C,G-J). ns, not significant, \**P*< 0.05; \*\**P* < 0.01; \*\*\**P* < 0.001. Also see Figure S5.

In the *in vitro* suppression assay, addition of neutralizing anti-IFN-γ antibody partially recovered the impaired suppressive activity of *IkE5*-deficient Treg cells (Figure 5E). Moreover, in *Rag2*^−/−^ mice co-transferred with *IkE5*-deficient Treg cells and naive CD4^+^CD25^−^CD45RB^hi^ T cells, neutralizing anti-IFN-γ antibody treatment prevented body weight loss, reduction of colon length, and massive infiltration of leukocytes into the colon (Figures 5F-5H). Consistent with these observations, administration of neutralizing anti-IFN-γ antibody abrogated the increase in the frequency of exTreg cells as well as the number of responder CD4^+^ T cells in *Rag2*^−/−^ mice receiving *IkE5*-deficient Treg cells (Figures 5I and 5J). Collectively, these results indicate that the overproduction of IFN-γ by *IkE5*-deletion is responsible for the impaired functional stability and suppressive function of Treg cells.

### The Foxp3-Ikzf1 complex controls chromatin architecture through the NuRD complex to repress gene expression in Treg cells

Since Ikzf1 associates with Foxp3 via IkE5 in Treg cells (Figure S1B), we first investigated whether Foxp3 and Ikzf1 bound to similar genomic sites in Treg cells by performing chromatin immunoprecipitation-sequencing (ChIP-seq). ChIP-seq analysis revealed that genomic distribution of both Foxp3 and Ikzf1 peaks were globally similar (Figures S6A); 20,583 and 22,275 peaks were identified as the sites with significant binding of Foxp3 and Ikzf1, respectively, in Treg cells, with more than 75% of Foxp3 peaks overlapped with Ikzf1 peaks (Figures S6B). The results suggested that Foxp3 and Ikzf1 shared a large number of target genes in Treg cells.

Next, to address whether genome-wide occupancy of Foxp3 would be affected by *IkE5*-deletion, we conducted ChIP-seq for Foxp3 in *IkE5*-deficient Treg cells. The assay revealed that 12.1% of Foxp3 peaks found in wild-type Treg cells were markedly reduced in intensity in *IkE5*-deficient Treg cells (designated Reduced Foxp3-binding Sites); 19.3% of Foxp3 peaks found in *IkE5*-deficient Treg cells were markedly enhanced compared with wild-type Treg cells (Enhanced Foxp3-binding Sites); the rest of Foxp3-binding sites were not altered in wild-type and *IkE5*-deficient Treg cells (Maintained Foxp3-binding Sites) (Figure 6A). The genomic distribution of these sites revealed that the Reduced and Enhanced Foxp3-binding Sites were much less localized at the promoter regions compared to the Maintained Foxp3-binding Sites, suggesting that the changes in Foxp3 binding due to *IkE5*-deletion preferentially occurred at the enhancer (intergenic and intron) regions (Figure 6B). Furthermore, 63.0% of the Reduced Foxp3-binding Sites in wild-type Treg cells located at the same sites of Ikzf1-binding, while 28.3% of the Enhanced Foxp3-binding Sites in *IkE5*-deficient Treg cells were bound by Ikzf1, suggesting that a sizable fraction of these Reduced or Enhanced Foxp3-bindings Sites are dependent on the presence of Ikzf1 for optimal Foxp3-binding (Figure 6C).

**Figure 6.**
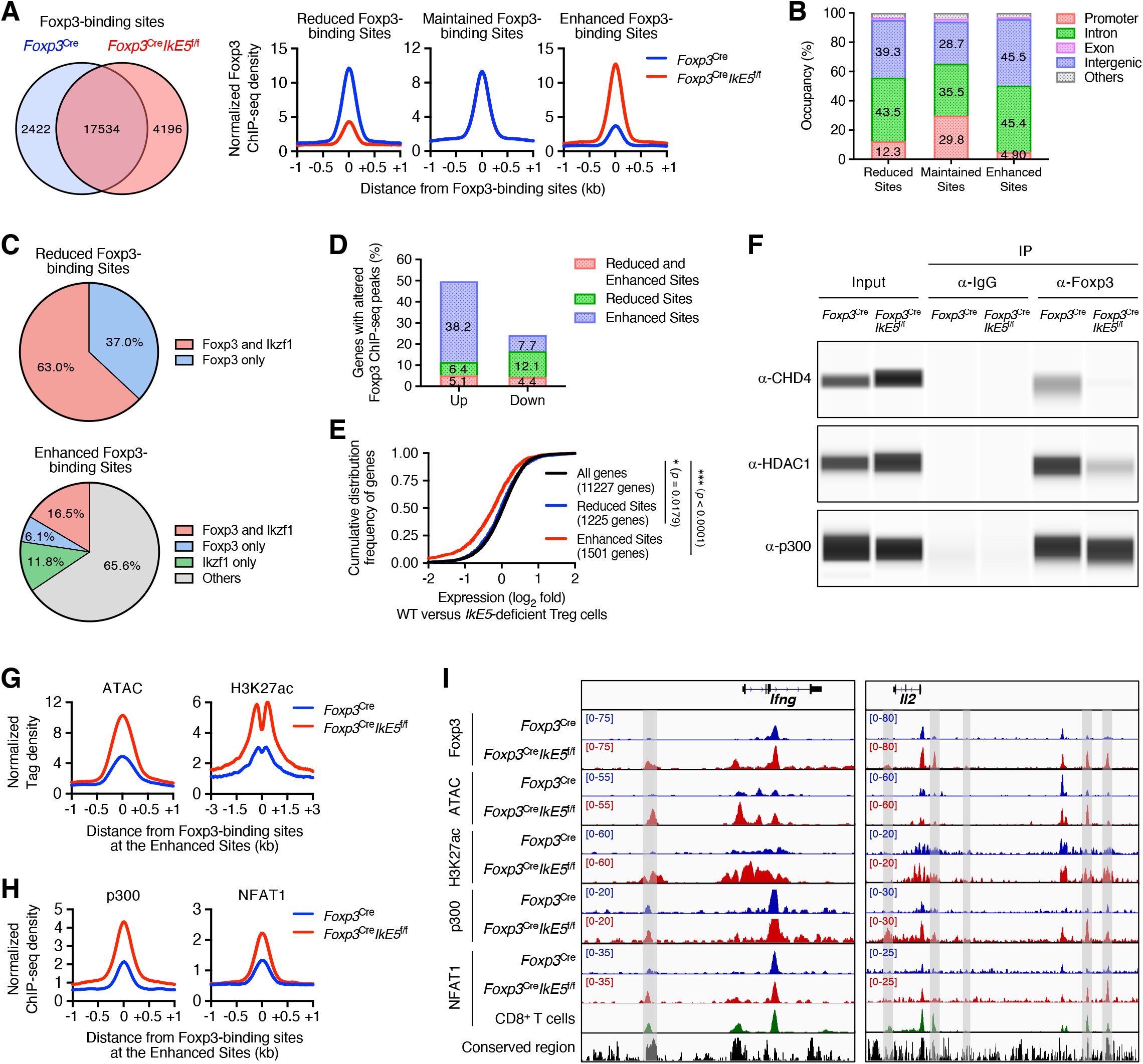
The Foxp3-Ikzf1 complex controls chromatin architecture through the NuRD complex to repress gene expression in Treg cells. (A) Venn diagram of Foxp3 ChIP-seq peaks in CD4^+^YFP^+^ Treg cells from *Foxp3*^Cre^ (blue) and *Foxp3*^Cre^*IkE5*^f/f^ (red) mice (left). Normalized density plots of Foxp3 binding peaks around the Reduced Foxp3-binding Sites, Maintained Foxp3-binding Sites and Enhanced Foxp3-binding Sites in CD4^+^YFP^+^ Treg cells from *Foxp3*^Cre^ (blue) and *Foxp3*^Cre^*IkE5*^f/f^ (red) mice (right). Normalized signal density is plotted within a window ± 1 kb centered on Foxp3-binding sites. (B) Peak annotation of the Reduced Foxp3-binding Sites, Maintained Foxp3-binding Sites and Enhanced Foxp3-binding Sites. (C) Pie chart illustrated the percentage of Foxp3- and Ikzf1-binding within the Reduced Foxp3-binding Sites (top) and Enhanced Foxp3-binding Sites (bottom). (D) Bar graph illustrated the percentage of genes harbored with altered Foxp3 binding sites among differentially Up and Down genes in *IkE5*-deficient Treg cells. (E) CDF analysis of the gene expression in wild-type and *IkE5*-deficient Treg cells for the genes corresponding to each the Reduced Foxp3-binding Sites and Enhanced Foxp3-binding Sites. (F) A representative image of interaction between Foxp3 and indicated factors, such as CHD4, HDAC1 and p300, in CD4^+^YFP^+^ Treg cells from *Foxp3*^Cre^ and *Foxp3*^Cre^*IkE5*^f/f^ mice by Co-IP. (G) Normalized density plots of ATAC and H3K27ac peaks around the Enhanced Foxp3-binding Sites in CD4^+^YFP^+^ Treg cells from *Foxp3*^Cre^ (blue) and *Foxp3*^Cre^*IkE5*^f/f^ (red) mice. Normalized signal density is plotted within a window ± 1-3 kb centered on Foxp3-binding sites. (H) Normalized density plots of indicated factors binding peaks around the Enhanced Foxp3-binding Sites in CD4^+^YFP^+^ Treg cells from *Foxp3*^Cre^ (blue) and *Foxp3*^Cre^*IkE5*^f/f^ (red) mice. Normalized signal density is plotted within a window ± 1 kb centered on Foxp3-binding sites. (I) Foxp3, p300, NFAT1, H3K27ac ChIP-seq and ATAC-seq signal tracks at the Treg down-regulated genes, such as *Ifng* and *Il2* genes loci in CD8^+^ T cells (green) as well as in CD4^+^YFP^+^ Treg cells from *Foxp3*^Cre^ (blue) and *Foxp3*^Cre^*IkE5*^f/f^ (red) mice. Data of CD8^+^ T cells (green) is from the previous report.^41^ Sequence conservation among vertebrates (black) is also shown. The Increased sites were highlighted in gray. Data are representative of two independent experiments (A-J). Also see Figure S6.

To determine then functional relevance of the alteration in Foxp3 binding, we integrated RNA-seq results and assessed the percentage of the genes possessing the altered Foxp3-binding sites among differentially Up- or Down-genes in *IkE5*-deficient Treg cells. Compared with 7.7% of Down-genes, 38.2% of Up-genes possessed the Enhanced Foxp3-binding Sites, implying a predominant contribution of the sites to the expression of Up-genes (Figure 6D). In agreement with these results, a CDF analysis revealed that the expression of the genes possessing the Enhanced Foxp3-binding Sites were markedly up-regulated in *IkE5*-deficient Treg cells (Figure 6E). These results collectively demonstrated that Ikzf1 association with Foxp3 was required for Treg-type gene expression, in particular, repression.

Foxp3 and its partners are known to form a large protein complex involving other TFs or epigenetic regulators for controlling the expression of target genes.^9^ We therefore examined by Co-IP analysis whether the components of the Foxp3 complexes were altered in *IkE5*-deficient Treg cells. Notably, core members of the nucleosome remodeling and deacetylase (NuRD) complex, CHD4 and HDAC1, were dissociated from Foxp3 complex by *IkE5*-deficiency (Figure 6F). Next, since the NuRD complex is a chromatin remodeling complex having nucleosome remodeling and histone deacetylase activities,^40^ we examined whether the Foxp3-Ikzf1 complex might suppress the expression of target genes by regulating chromatin accessibility and histone acetylation trough the NuRD complex. Assay for transposase-accessible chromatin (ATAC)-seq and histone H3 on lysine 27 acetylation (H3K27ac) ChIP-seq analyses indeed revealed that chromatin accessibility and enhancer activity at the Enhanced Foxp3-binding Sites were low in wild-type Treg cells compared to other sites (Figure S6C), but these activities were strongly augmented by *IkE5*-deficiency (Figure 6G). Consistent with these results, the histone acetyl transferase (HAT), p300, interacted with Foxp3 and its binding to the Enhanced Foxp3-binding Sites were augmented in *IkE5*-deficient Treg cells (Figures 6F and 6H). Because Foxp3 complexes are heterogeneous in composition,^9^ these results indicate that Foxp3 may form a repressive complex with Ikzf1 and compete with the active Foxp3-p300 complex in controlling gene transcription.

Motif enrichment analysis also revealed that NFAT motif was mostly enriched in the Enhanced Foxp3-binding Sites (Figure S6D). Using a published dataset,^41^ we further found that 31.5% of the Enhanced Foxp3-binding Sites bound NFAT1 in CD8 T cells, compared to only 2.5% in wild-type Treg cells (Figure S6E). NFAT1-binding to the Enhanced Foxp3-binding Sites were indeed increased in *IkE5*-deficient Treg cells (Figure 6H), suggesting an inhibition of NFAT1-binding to the Enhanced Foxp3-binding Sites by the Foxp3-Ikzf1 complex. We further confirmed that epigenetic signatures as well as epigenetic regulators binding were altered at the Enhanced Foxp3-binding Sites around the Treg down-regulated genes loci, including *Ifng* and *Il2* in *IkE5*-deficient Treg cells (Figure 6I). In contrast, these alterations were not detected at the Treg up-regulated genes loci, including *Ctla4*, *Il2ra* and *Tnfrsf18* (Figure S6F).

Collectively, these findings indicate that the Foxp3-Ikzf1 complex competes with co-activators such as p300 and NFAT1 by inducing closed chromatin architecture through the NuRD complex in an Ikzf1-dependent manner, consequently suppressing the expression of its target genes.

### Both Ikzf1 and Ikzf3 association with Foxp3 is required for the maintenance of Treg cell homeostasis in mice and humans

Recently, Carolina et al. reported that *dLcK*^Cre^*Ikzf1*^f/f^ mice, in which Ikzf1 was specifically deleted in mature T cells, showed a normal frequency of splenic Treg cells and no overt spontaneous autoimmunity during 3 months’ observation,^42^ suggesting possible defects other than the Ikzf1 anomaly in *Foxp3*^Cre^*IkE5*^f/f^ mice. Ikzf1 functions by forming not only the homodimer but also a heterodimer with other Ikzf family members,^22^ suggesting that *IkE5*-deficiency might affect function of its heterodimeric partners in Treg cells. To identify putative Ikzf family proteins that contribute to the phenotypes of *IkE5*-deficient Treg cells, we generated Treg cells deficient in each Ikzf family member, or in combination, by *in vitro* CRISPR/Cas9-mediated editing (ICE). The ICE system efficiently and specifically disrupted the targeted Ikzf family proteins in primary Treg cells (Figure S7A). In these conditions, double-deficiency of Ikzf1 and Ikzf3 significantly induced IFN-γ production and generated exTreg cells compared to control, to a similar extent as quadruple-deficiency of Ikzf1, Ikzf2, Ikzf3 and Ikzf4 (Figure 7A). On the other hand, double-deficiency of Ikzf2 and Ikzf4 as well as single-deficiency of each Ikzf family protein failed to induce IFN-γ production and exTreg cell generation (Figure 7A). These results suggest that the *IkE5*-deletion affects not only Ikzf1 but also Ikzf3 function in Treg cells.

**Figure 7.**
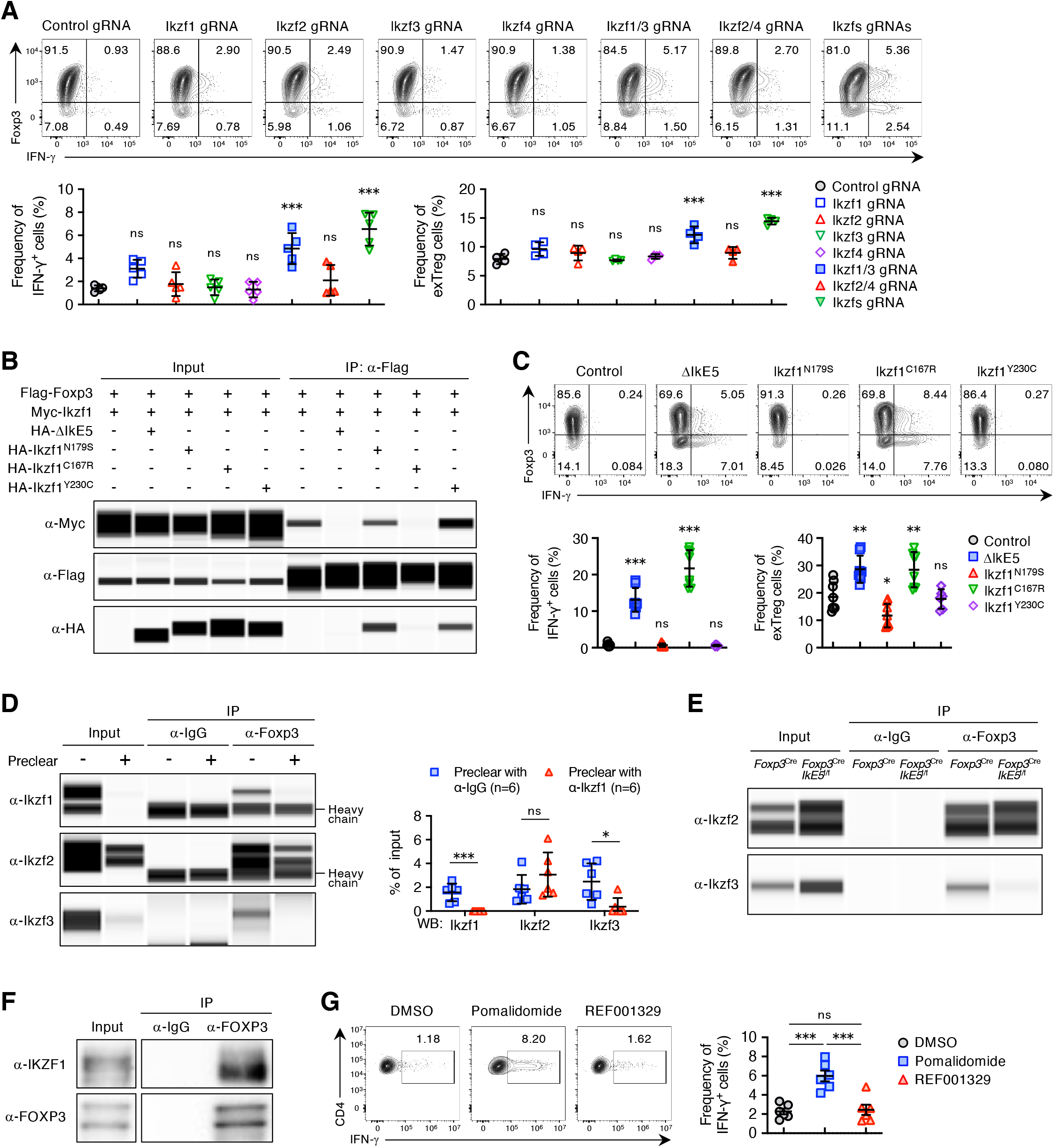
Both Ikzf1 and Ikzf3 association with Foxp3 is required for the maintenance of Treg cell homeostasis in mice and humans. (A) Cas9 protein and indicated gRNAs were introduced into the activated Treg cells (2 × 10^6^) by nucleofection. Nucleofected cells were stimulated with Dynabeads mouse CD3/CD28 T cell stimulator (25 μl/ml) and IL-2 (1500 U/ml) for 3 days, followed by flow cytometry analysis. A representative flow cytometry plot of CD4^+^ T cells (up) and frequencies of CD4^+^IFN-γ^+^ cells (bottom left) and exTreg cells (bottom right) in nucleofected cells (mean ± SD, *n* = 4-5 per group). (B) A representative image of the interaction between Foxp3 and Ikzf1 (WT) in the presence of Ikzf1-mutants in HEK293T cells by Co-IP. (C) Purified CD4^+^FYP^+^ Treg cells (1 × 10^5^) were stimulated with α-CD3 (1 μg/ml), α-CD28 (1 μg/ml) and IL-2 (100 U/ml) for 24 hrs. Activated Treg cells were transduced with fresh retrovirus supernatant by spin-infection, and then were cultured under the above conditions and harvested on day 4 for flow cytometric analysis. A representative flow cytometry plot of CD4^+^ T cells (up) and frequencies of CD4^+^IFN-γ^+^ cells (bottom left) and exTreg cells (bottom right) in transduced cells (mean ± SD, *n* = 6-8 per group). (D) Whole cell lysates from CD4^+^FYP^+^ Treg cells were pre-cleared with 5 μg ant-IgG (−) or anti-Ikzf1 (+) antibody overnight at 4℃. Subsequently, pre-cleared lysates were immunoprecipitated with 5 μg ant-IgG or anti-Foxp3 antibody overnight at 4℃, followed by Simple Western assay with anti-Ikzf1, anti-Ikzf2 and anti-Ikzf3 as primary antibodies (WB). A representative image of Simple Western (left) and percentages of immunoprecipitates relative to amount in input from pre-cleared lysates (right)(mean ± SD, *n* = 6 per group). (E) A representative image of interaction between Foxp3 and indicated factors, such as Ikzf2 and Ikzf3, in CD4^+^YFP^+^ Treg cells from *Foxp3*^Cre^ and *Foxp3*^Cre^*IkE5*^f/f^ mice by Co-IP. (F) A representative image of interaction between FOXP3 and IKZF1 in human CD4^+^CD25^+^CD127^lo^ naive Treg cells by Co-IP. (G) Sorted human CD4^+^CD25^+^CD45RA^+^ and CD4^+^CD25^hi^CD45RA^−^ Treg cells (5 × 10^4^) were cultured with Dynabeads Human CD3/CD28 T cell stimulator and IL-2 (100 U/ml) in the presence of DMSO, Pomalidomide (10 μM) or REF001329 (10 μM) for 9 days, followed by flow cytometry analysis. A representative flow cytometry plot of CD4^+^Foxp3^+^ T cells (up) and frequencies of CD4^+^Foxp3^+^IFN-γ^+^ Treg cells (bottom) in cultured cells (mean ± SEM, *n* = 6 per group). Data are summary of at least three independent experiments (A,C,D,G) or representative of three independent experiments (B,E,F). *P* values determined by ordinary one-way ANOVA followed by Dunnett’s multiple comparisons (A,C) or unpaired *t*-tests followed by Holm-Šídák multiple comparisons test (D) or ordinary one-way ANOVA followed by Tukey’s multiple comparisons test (G). ns, not significant, \**P*< 0.05; \*\**P*< 0.01; \*\*\**P* < 0.001. Also see Figure S7.

The IkE5-deleted Ikzf1 mutant (ΔIkE5) has been reported to exert a dominant-negative effect in B cells.^30^ Moreover, various germline IKZF1 point mutations, which show four different mechanisms of action; 1) haploinsufficiency mutations, 2) dimerization defective mutations, 3) dominant-negative mutations, and 4) gain-of-function mutations, were recently identified in patients with immunodeficiency and autoimmune diseases.^27^ To clarify whether the effect of *IkE5*-deficiency in Treg cells was due to dominant-negative effect of ΔIkE5 or its dissociation from Foxp3, we conducted overexpression of several Ikzf1-mutants in Treg cells. Similar to *IkE5*-deficient Treg cells, overexpression of ΔIkE5 led to inhibition of interaction between Foxp3 and Ikzf1, and promoted IFN-γ production and exTreg cell generation in Treg cells (Figure 7B and 7C). In contrast, overexpression of a dominant-negative mutant Ikzf1^N179S^ (N159S in humans) failed to inhibit above interaction and to induce Treg cell instability (Figure 7B and 7C). Of note, a haploinsufficiency mutant Ikzf1^C167R^ (C147R in humans) that inhibited the interaction between Foxp3 and Ikzf1 promoted IFN-γ production and exTreg cell generation, while another haploinsufficiency mutant Ikzf1^Y230C^ (Y210C in humans) that had no inhibitory effect on their interaction maintained Treg cell stability (Figure 7B and 7C). These results collectively suggest that dissociation of Ikzf1 from Foxp3 by *IkE5*-deficiency is causative of functional instability in *IkE5*-deficient Treg cells.

Kwon *et al*. previously demonstrated that Ikzf2 and Ikzf3 each form different Foxp3 complexes in Treg cells,^20^ which suggested to us that Ikzf1 might belong to the same Foxp3 complex with Ikzf3 in Treg cells. To address this possibility, we performed a pre-cleared Co-IP experiments in which Treg cell lysates were pre-cleared with anti-Ikzf1 or anti-IgG antibody before immunoprecipitation of Foxp3. Pre-clearing the lysates with anti-Ikzf1 antibody significantly depleted Ikzf3-Foxp3 complexes, while it did not affect Ikzf2-Foxp3 complexes (Figure 7D). Moreover, we confirmed the dissociation of Ikzf3, but not Ikzf2, from Foxp3 in *IkE5*-deficient Treg cells (Figure 7E). These observations collectively indicate that Ikzf1 and Ikzf3 belong to the same Foxp3 complex in Treg cells, while Ikzf2 belongs to a different one, which explains the functional defect of both Ikzf1 and Ikzf3 by *IkE5*-deletion. The results also indicate that both Ikzf1 and Ikzf3 association with Foxp3 is required for the maintenance of Treg cell homeostasis.

To address whether IKZF1 would similarly associate with FOXP3 in humans, we conducted Co-IP analysis with peripheral blood CD4^+^CD25^+^CD127^lo^ naive Treg cells from healthy donors, and confirmed the interaction between IKZF1 and FOXP3 (Figure 7F). We next evaluated the role of IKZF family proteins in human Treg cells using a protein knock-down strategy with small molecules. Thalidomide analogues, such as Lenalidomide and Pomalidomide, can induce degradation of IKZF1 and IKZF3 by recruiting them to the CRL4(CRBN) E3 ubiquitin ligase.^43–46^ Moreover, one compound, 3-(5-(1-(4-(difluoromethoxy)benzyl)piperidin-4-yl)-1-oxoisoindolin-2-yl)piperidine-2,6-dione, has recently been reported as a potent and selective degrader for IKZF2 and IKZF4 in the International Patent Applications (WO 2019/038717 AI). Therefore, we synthesized the compound (designated REF001329) and *in vitro* cultured human Treg cells with Pomalidomide or REF001329 in the presence of anti-CD3, anti-CD28 and IL-2. IKZF1 and IKZF3 or IKZF2 and IKZF4 were indeed selectively degraded by the treatment with Pomalidomide or REF001329, respectively (Figure S7B). Under these conditions, overproduction of IFN-γ was observed in Pomalidomide-treated Treg cells, but not in those treated with REF001329 (Figure 7G), indicating that both IKZF1 and IKZF3 are required for the suppression of IFN-γ in human Treg cells. These findings taken together suggest that Ikzf1 and Ikzf3 cooperatively play crucial roles in the maintenance of Treg cell homeostasis in mice and humans.

## DISCUSSION

Foxp3 forms the molecular complexes with numerous factors that facilitate epigenetic remodeling through regulation of histone modifications and DNA methylation, thereby activating or repressing target genes. For example, Foxp3 is thought to induce histone H3 acetylation near promoters and enhancers of its target genes, such as CD25, CTLA4, and GITR, by associating with p300 and other key transcriptional co-activators, leading to the activation of target genes.^10,15^ In contrast, when Foxp3 functions as a repressor, Foxp3 may silence the expression of target genes such as IL-2 and IFN-γ by inducing deacetylation of histone H3 via recruiting histone deacetylase and transcriptional co-repressors, including histone deacetylase 3 (HDAC3), carboxy-terminal binding protein 1 (CtBp1) and nuclear receptor co-repressors (NCoR1/NCoR2).^10,23,47^ This report has shown another inhibitory mechanism mediated by an association of Foxp3 with Ikzf1.

Ikzf1 is an essential regulator of lymphocyte differentiation and functions primarily as a gene repressor by interacting with the NuRD complex.^48–50^ Our results indicate that Foxp3 forms a repressive complex with the NuRD complex in an Ikzf1-dependent manner and suppresses the expression of its target genes by competing directly with co-activators, such as p300 and NFAT1, through induction of closed chromatin architecture. Consistent with our findings, many studies have suggested a direct gene regulation model for Foxp3 in determining Treg cell identity.^10–12,16,18^ On the other hand, a recent study has shown that Foxp3 augments the Treg-type chromatin accessibility largely in an indirect manner by tuning the activity of other chromatin remodeling TFs such as TCF1.^25^ Hence, it remains debatable whether Foxp3 regulates Treg-specific transcription directly or indirectly by tuning intermediates. Interestingly, our RNA-seq analysis has shown that the expression of several TFs, such as *Bhlhe40* and *Nfat2*, are significantly up-regulated in *IkE5*-deficient Treg cells. Bhlhe40 and NFAT2 are key regulators of T-cell activation and cytokine production.^51,52^ These results taken together suggest that the Foxp3-Ikzf1 complex may also indirectly suppress the expression of target genes through repression of transcriptional regulators in Treg cells.

As a model for Foxp3-mediated regulation of gene expression, Kwon *et al*. have shown that Foxp3 exists in distinct multimolecular complexes which can be segregated into two groups based on co-factors, function and nuclear localization.^20^ Specifically, Foxp3 actively regulates the expression of its target genes, both positively and negatively, when complexed with RELA, IKZF2 and KAT5, and localizes to the center of the nucleus. In contrast, Foxp3 is inactive when complexed with YY1, IKZF3 and EZH2, has a diminished activity in both the activation and repression of its target genes, and is sequestrated in the periphery of the nucleus.^20^ We have clarified in the present study that Ikzf1 forms Foxp3 complex with Ikzf3, one of the components of the inactive Foxp3 complex in Kwon’s model, in activated Treg cells, and that both Ikzf1 and Ikzf3 play an important role in the suppression of IFN-γ production and exTreg generation from Treg cells. In addition, DuPage et al. has reported that EZH2, another component of the inactive Foxp3 complex in Kwon’s model, is required for stabilization of Foxp3-driven transcriptional program in activated Treg cells.^53^ To explain these discrepancies, it should be noted that Kwon *et al*. mainly used Foxp3-transduced T cells in their study.^20^ It has been shown that Foxp3-transduced T cells exhibit a transcriptional program and a CpG hypomethylation pattern distinct from those in endogenous naturally occurring Treg cells,^13,54^ and that Foxp3 becomes functional and engages in gene repression after TCR stimulation.^14^ Since we and DuPage et al. analyzed natural Treg cells activated with anti-CD3 and anti-CD28, it is likely that the inactive Foxp3 complex becomes functional upon TCR stimulation.

The Ikzf1 isoforms lacking ZFs of the N-terminus have been reported to exert a dominant-negative effect by forming dimers with other Ikzf family members and interfering with their DNA-binding activity.^22^ Since *IkE5*-deletion generates an Ikaros isoform lacking the two N-terminal ZFs, it is predicted to have a dominant-negative effect. Joshi et al. indeed showed that *IkE5*-deficient mice exhibited a dominant-negative phenotype in B cells.^30^ We have, however, experimentally demonstrated that instability of *IkE5*-deficient Treg cells was not due to a dominant-negative effect by ΔIkE5 since overexpression of a dominant-negative mutant Ikzf1^N179S^ could not induce IFN-γ production and exTreg generation in Treg cells. Furthermore, in contrast to impaired IL-2-mediated activation of STAT5 in *Ikzf2*-deficient Treg cells,^24^ IL-2-STAT5 signaling was not affected in *IkE5*-deficient Treg cells, suggesting that Ikzf2 function is intact in the latter. These results support our finding that ΔIkE5 does not exert a dominant-negative effect in Treg cells. On the other hand, dissociation of Ikzf1 from Foxp3 by *IkE5*-deletion can be a main cause for instability of *IkE5*-deficient Treg cells because overexpression of ΔIkE5 or a haploinsufficient Ikzf1^C167R^ mutation, which inhibited the interaction between Foxp3 and Ikzf1, similarly induced IFN-γ production in Treg cells and generated exTreg cells. In addition, patients carrying IKZF1 C147R mutation (C167R in mice) reportedly show a reduction of Treg cells and development of autoimmune disease.^28^ However, with the differences in the phenotype, such as serum Ig levels and IFN-γ production, between our *IkE5*-deficient mouse model and the patients with IKZF1 germline mutations, further investigation of Ikzf1 mutations, their functional anomalies, and disease phenotype is necessary to elucidate the function of Ikzf1 in Treg and other immune cells.

We have found that pomalidomide treatment compromises Treg cell homeostasis with a decrease in IKZF1 and IKZF3 in human Treg cells, as observed with Ikzf1 and Ikzf3 double-deficiency in mice. Immunomodulatory drugs (IMiDs) such as lenalidomide and pomalidomide have been used in the treatment of cancers including multiple myeloma and myelodysplastic syndrome, and an inflammatory skin pathology associated with Hansen’s disease because of their anti-tumor, anti-angiogenic and anti-inflammatory properties.^55^ Furthermore, IMiDs are promising for the treatment of autoimmune disorders such as systemic lupus erythematosus and inflammatory bowel disease.^56,57^ Recent studies have revealed that cereblon, an ubiquitin E3 ligase, is a receptor of IMiDs, and that both IKZF1 and IKZF3 are target molecules of IMiDs.^43,44^ Specifically, IMiDs have been reported to induce cytotoxicity in multiple myeloma and MDS del(5q) cell lines by a cereblon-dependent degradation of IKZF1 and IKZF3 as one of the mechanisms of their anti-tumor activity.^43,44^ moreover, IMiDs may contribute to anti-inflammatory effects by promoting Treg cell survival and suppressing pathogenic Th17 cell differentiation via increased IL-2 production from activated T cells through degradation of IKZF1 and IKZF3.^58,59^ To date, however, these proposals on anti-inflammatory property of IMiDs have lacked sufficient experimental support or large-scale controlled clinical trials. Our findings thus provide not only a new mechanism of the anti-cancer activity of IMiDs, but also clues to explaining their anti-inflammatory effects for treating immunological diseases.

In conclusion, the present study demonstrates that the Foxp3/Ikzf1/Ikzf3 complex exerts gene-repressing function via chromatin remodeling at the target gene loci in Treg cells. Our findings can be exploited to devise novel immunotherapies of immunological diseases and cancer.

## METHODS

### Mice

*IkE5*^f/f^ mice^30^ were crossed with *Foxp3*^Cre^ transgenic mice^31^ or *Foxp3*^eGFP-Cre-ERT2^ mice^36^ to generate *Foxp3*^Cre^*IkE5*^f/f^ mice or *Foxp3*^eGFP-Cre-ERT2^*IkE5*^f/f^ mice, respectively. *Foxp3*^Cre^*IkE5*^f/f^ mice and *Rosa26*^rfp^ reporter mice^33^ were bred to yield *Foxp3*^Cre/+^*IkE5*^f/f^*R26*^rfp/+^ mice. *Foxp3*^Cre/+^*IkE5*^f/f^*R26*^rfp/+^ mice were crossed with *Ifng^−/−^* mice to generate *Foxp3*^Cre/+^*IkE5*^f/f^*Ifng^−/−^R26*^rfp/+^ mice. CD45.1^+^C57BL/6 and CD45.1^+^*Rag2*^−/−^ mice were bred in our animal facility. The animal experiments except murine tumor models were performed by using age-matched, 3 to 4-week-old mice. *Foxp3*^eGFP-Cre-ERT2^*IkE5*^f/f^ and *Rag2*^−/−^ recipient mice were used at 8 to 12 weeks of age for experiments. All mice used were maintained under specific pathogen-free conditions and all experiments were performed in accordance with guidelines for animal welfare set by Osaka University.

### Plasmids construction

Murine Ikzf1 and Foxp3 cDNA were amplified by RT-PCR, and inserted into pCMV-Tag vectors (Agilent) to generate Tag fused-Ikzf1 and -Foxp3. The deletion fragments and point-mutants of Ikzf1 were amplified by RT-PCR, and inserted into pCMV-Tag vectors (Agilent) and MSCV-NGFR vector to generate Tag fused-Ikzf1 mutants and retroviral Ikzf1 mutants, respectively.

### Cell preparation and culture conditions

Mouse Treg cell sorting from peripheral lymphoid organs was performed as previously described.^60^ Briefly, single cell suspensions were prepared from spleen and lymph nodes, and were pre-enrichment of CD4^+^ cells using a CD4^+^ T cell isolation kit, mouse (Miltenyi Biotec). Enriched cells were stained with Live/Dead cell viability dye and antibodies for surface markers as follows, CD4, CD25, CD8, CD3, CD44, CD62L and CD45RB. Then, CD4^+^CD8^−^FYP^+^ cells were sorted as Treg cells by FACSAria II (BD Biosciences). In some experiments, CD4^+^CD25^−^YFP^−^CD44^lo^CD62L^hi^ cells or CD4^+^CD25^−^CD45RB^hi^ cells were prepared as naive CD4^+^ T cells. CD3^−^CD25^−^YFP^−^ cells from spleen were prepared as CD3^+^ T cell-depleted splenocytes. For mouse cell culture, we used RPMI 1640 (Nacalai tesque) supplemented with 10% fetal calf serum (FCS)(v/v), 60 μg/ml penicillin G (Nacalai tesque), 100 μg/ml streptomycin (Nacalai tesque) and 0.1 mM 2-mercaptoethanol (Thermo Fisher Scientific).

For purification of human Treg cells, human CD4^+^ T cells were purified from frozen peripheral blood mono-nuclear cells (PBMCs) (STEMCELL Technologies) with EasySep Human CD4^+^ T Cell Isolation Kit (STEMCELL Technologies). Enriched CD4^+^ cells were stained for 30 min on ice with antibodies against anti-CD4, anti-CD25 and anti-CD45RA antibodies. CD4^+^CD25^+^CD45RA^+^ and CD4^+^CD25^hi^CD45RA^−^ cells were sorted by MA900 cell sorter (SONY Biotechnology). For Co-IP experiment, naive Treg cells were purified by using EasySep Human CD4^+^CD127^low^CD25^+^ Regulatory T Cell Isolation Kit (STEMCELL Technologies). For human cell culture, we used RPMI 1640 (Thermo Fisher Scientific) supplemented with 10% fetal calf serum (FCS)(v/v)(Thermo Fisher Scientific), Penicillin-Streptomycin Solution (x100) (FUJIFILM Wako). The present study was approved by the institutional ethics committees of Osaka University.

### Flow cytometry analysis

Flow cytometry analysis was performed as previously described.^61^ Cells were first incubated with anti-CD16/32 then stained with Live/Dead cell viability dye, and antibodies for surface markers. Cells were subsequently fixed and permeabilized with a Foxp3/Transcription Factor Staining Buffer Set (eBioscence) according to the manufacturer’s instructions. For intracellular cytokine staining, cells were incubated with Cell Stimulation Cocktail (plus protein transport inhibitors) (eBioscience) for 4 hrs or Cell Activation Cocktail (Biolegend) in the presence of Brefeldin A and Monensin for 5 hrs before staining.

For phosphorylated STAT5 staining, cells were first starved for 45 min at 37°C, followed by stimulation with IL-2 (100 U/ml) for 30 min at 37°C. Stimulated cells were then subjected to BD Phosflow Lyse/Fix Buffer and BD Phosflow Perm Buffer III (BD Biosciences), according to the manufacturer’s instructions. Data were acquired through a FACSCanto II (BD Biosciences) for mouse samples or a CytoFLEX LX (Beckman Coulter) for human samples, and analyzed with FlowJo software (Tree Star Inc).

### Immunoblot analysis and Simple western assay

Immunoblot analysis was performed as described previously.^62^ Briefly, cells were washed, lysed in sample buffer, and boiled. Whole cell lysates were separated by SDS-PAGE and transferred to a polyvinylidene difluoride membrane by using iBlot 2 Dry Blotting System (Thermo Fisher Scientific). The membrane was blocked with StartingBlock (TBS) Blocking Buffer (Thermo Fisher Scientific) and probed with the following primary antibodies: anti-IKZF1 (Cell Signaling Technology), anti-IKZF2 (Cell Signaling Technology), anti-IKZF3 (Cell Signaling Technology), anti-IKZF4 (Novus Biologicals), anti-FOXP3 (Cell Signaling Technology), anti-β-Actin (Invitrogen), anti-Flag-M2 (Sigma-Aldrich) or anti-c-Myc-Tag (Cell Signaling Technology). The membrane was then probed with appropriate secondary antibodies conjugated to HRP and visualized with SuperSignal West Pico PLUS Chemiluminescent Substrate (Thermo Fisher Scientific) according to the manufacturer’s instructions. Anti-β-Actin was used as a loading control.

For the Simple western assay, the Jess Simple Western System (ProteinSimple) is an automated capillary-based size separation and nano-immunoassay system. We followed the manufacturer’s standard method. Briefly, the total cell lysates were mixed with 0.1x sample buffer and Fluorescent 5x master mix (ProteinSimple) in the presence of fluorescent molecular weight markers and 400 mM dithiothreitol (ProteinSimple). The samples were separated in capillaries as they migrated through a separation matrix. A Protein Simple proprietary photoactivated capture chemistry was used to immobilize separated proteins on the capillaries. After a wash step, the primary antibodies: anti-Ikzf1 (Cell Signaling Technology), anti-Ikzf2 (Cell Signaling Technology), anti-Ikzf3 (Santa Cruz Biotechnology), anti-CHD4 (Sigma-Aldrich), anti-HDAC1 (Cell Signaling Technology), anti-p300 (Sigma-Aldrich) or anti-GAPDH (Cell Signaling Technology), and HRP-conjugated secondary antibodies (ProteinSimple): anti-rabbit secondary antibody or anti-mouse secondary antibody, were incubated for 50 min and 30 min, respectively. The chemiluminescent revelation was established with peroxyde/luminol-S (ProteinSimple). Digital image of chemiluminescence of the capillary was captured with Compass Simple Western software (ProteinSimple) that calculated automatically heights (Chemiluminescence intensity), area, and signal/noise ratio. If no signal of target factors was calculated, band intensity was defined as 0. Results could be visualized as electropherograms representing peak of chemiluminescence intensity and as lane view from signal of chemiluminescence detected in the capillary. An internal system control was included in each run.

### Co-immunoprecipitation (Co-IP)

Expression vectors encoding Foxp3 and Ikzf1 and/or Ikzf1 mutants were transfected into HEK293T cells (2 × 10^5^) with FuGENE HD (Promega). 48h after transfection, cells were harvested and lysed in IP lysis buffer (Thermo Fisher Scientific). Immunoprecipitation was performed using 2 μg anti-Flag-M2 antibody (Sigma-Aldrich) in the presence of DynaBeads IgG magnetic beads (25 μl/sample)(Thermo Fisher Scientific) overnight at 4℃. Equivalent amounts of protein from whole cell lysates (Input) or immunoprecipitates (IP) were analyzed by Immunoblot analysis.

In primary mouse Treg cells, purified CD4^+^FYP^+^ Treg cells (1 × 10^5^) were cultured with Dynabeads mouse CD3/CD28 T cell stimulator (25 μl/ml) (Thermo Fisher Scientific) and IL-2 (1500 U/ml) for 7 days. Activated Treg cells were then stimulated with Cell Stimulation Cocktail (eBioscience) for 1 hr at 37 ℃ and lysed in IP lysis buffer (Thermo Fisher Scientific). Immunoprecipitation was performed with 5 μg anti-IgG (Sigma-Aldrich) or 5 μg anti-Foxp3 (eBioscience) antibody in the presence of DynaBeads IgG magnetic beads (50 μl/sample) (Thermo Fisher Scientific) overnight at 4℃. Equivalent amounts of protein from whole cell lysates (Input) or immunoprecipitates (IP) were analyzed by Simple Western assay.

In human Treg cells, purified CD4^+^CD25^+^CD127^lo^ naive Treg cells (5 × 10^4^) were stimulated with Dynabeads Human T-activator CD3/CD28 (12.5 μl/ml) (Thermo Fisher Scientific) and IL-2 (100 U/ml) for 9 days. Expanded Treg cells were harvested and lysed in RIPA buffer (Nacalai tesque). Immunoprecipitation was performed using 6 μg anti-FOXP3 antibody (eBioscience) and Protein G Dynabeads (20 μl/sample) for 3 hrs. Equivalent amounts of protein from whole cell lysates (Input) or immunoprecipitates (IP) were analyzed by Immunoblot analysis.

### Pre-clearing Co-IP

Purified CD4^+^FYP^+^ Treg cells (1 × 10^5^) were cultured with Dynabeads mouse CD3/CD28 T cell stimulator (25 μl/ml) (Thermo Fisher Scientific) and IL-2 (1500 U/ml) for 7 days, and lysed in IP lysis buffer (Thermo Fisher Scientific). Before immunoprecipitation, whole cell lysates were pre-cleared with 5 μg anti-IgG (Cell Signaling Technology) or 5 μg anti-Ikzf1 (Cell Signaling Technology) antibody in the presence of DynaBeads IgG magnetic beads (50 μl/sample) (Thermo Fisher Scientific) overnight at 4℃. Subsequently, pre-cleared lysate were immunoprecipitated with 5 μg ant-IgG (Sigma-Aldrich) or 5 μg anti-Foxp3 (eBioscience) antibody in the presence of DynaBeads IgG magnetic beads (50 μl/sample) (Thermo Fisher Scientific) overnight at 4℃. Equivalent amounts of protein from supernatants (Input) or immunoprecipitates (IP) were analyzed by Simple Western assay.

### Enzyme linked immunosorbent assay (ELISA)

Serum was collected from *Foxp3*^Cre^ and *Foxp3*^Cre^*IkE5*^f/f^ mice at 3 to 4 weeks of age and analyzed for concentration of IgG1, IgM and IgE with Mouse Uncoated ELISA kits (Invitrogen) according to the manufacturer’s instructions. Anti-double strand DNA (dsDNA) and anti-parietal cell (PC) antibodies in serum were respectively measured with a LBIS Mouse anti-dsDNA ELISA Kit (FUJIFILM Wako Shibayagi Corporation) and a Qualitative Mouse Gastric Parietal Cell Antibody (Anti-PC) ELISA Kit (MYBioSource), according to the manufacturer’s protocol. The absorbance was measured at 450 nm with Nivo multimode microplate reader (PerkinElmer).

### Histological analysis

Freshly-isolated tissues were immediately fixed by 4% Paraformaldehyde (Nacalai tesque). Hematoxylin and eosin (H&E) staining and microscopy slide preparation was performed by the Center for Anatomical, Pathological and Forensic Medical Research, Kyoto University Graduate School of Medicine. Stained sections were subjected to scoring of disease severity, in a double-blinded manner, based on the following criteria.

Dermatitis: 0, normal (no inflammation); 1, mild inflammation; 2, moderate inflammation; 3, marked inflammation (thickness and tissue destruction).

Gastritis: 0, no inflammation; 1, submucosal inflammation; 2, mild mucosal inflammation; 3, intermediate mucosal inflammation with destruction of gastric glands; 4, severe mucosal inflammation with loss of parietal cells.

Pneumonitis: 0, no inflammation; 1, mild inflammation; 2, intermediate inflammation; 3, severe inflammation and tissue destruction.

Hepatitis: 0, normal (no pathology); 1, mild (1-3 abnormal areas); 2, moderate (3-5 abnormal areas); 3, severe (> 5 abnormal areas).

Colitis: 0, no inflammation; 1, minimal scattered mucosal inflammatory cell infiltrates, with or without minimal epithelial hyperplasia; 2, mild scattered mucosal and submucosal inflammatory cell infiltrates with mild epithelial hyperplasia; 3, moderate scattered mucosal and submucosal inflammatory cell infiltrates with moderate epithelial hyperplasia and mucin depletion; 4, marked scattered mucosal and submucosal inflammatory cell infiltrates that were associated with ulceration with marked epithelial hyperplasia and mucin depletion; 5, marked transmural inflammation with severe ulceration and loss of intestinal glands.

### Treg cell stability assay

For the *in vitro* assay, purified CD4^+^FYP^+^ Treg cells (1 × 10^5^) were cultured with Dynabeads mouse CD3/CD28 T cell stimulator (25 μl/ml) (Thermo Fisher Scientific) and IL-2 (100 U/ml) in the presence of 10 μg/ml anti-IFN-γ (eBioscience) or 10 μg/ml anti-IgG antibody (eBioscience) for 7 days. Cultured cells were analyzed by flow cytometry after removing Dynabeads.

For the *ex vivo* assay, purified CD4^+^FYP^+^ Treg cells (1 × 10^5^) from *Foxp3*^Cre/+^, *Foxp3*^Cre/+^*IkE5*^f/f^ or *Foxp3*^Cre/+^*IkE5*^f/f^*Ifng*^−/−^ mice were intravenously transferred into *Rag2*^−/−^ recipients. At day 7-8 after transfer, peripheral lymphoid organs were collected from recipient mice and subjected to flow cytometric analysis.

For human Treg cells, purified Treg cells (5 × 10^4^) were cultured with Dynabeads Human T-activator CD3/CD28 (12.5 μl/ml) (Thermo Fisher Scientific) and IL-2 (100 U/ml) stimulations in the presence of DMSO, 10 μM Pomalidomide (Tokyo Chemical Industry) or 10 μM REF001329 (Fujii Memorial Research Institute) for 9 days, followed by flow cytometry analysis.

### CpG methylation analysis by bisulfite sequencing

Genome DNA was collected from purified Treg cells (1 × 10^5^) by phenol-chloroform extraction. 10-100 ng of genome DNA was subjected to bisulfite reaction using MethylEasy Xceed Rapid DNA Bisulfite Modification Kit (Human Genetic Signatures) following the manufacturer’s instruction. PCR primers, conditions, and methods for DNA sequencing are previously described.^13^

### *In vitro* suppression assay

Purified CD4^+^ naïve T (Tresp) cells from CD45.1^+^ C57BL/6 mice were stained with CellTrace violet (CTV) (Thermo Fisher Scientific). Tresp cells (5 × 10^4^) were cultured with CD3^+^ T cell-depleted splenocytes (1 × 10^5^) and CD4^+^FYP^+^ Treg cells at indicated Treg : Tresp ratios in the presence of 1 μg/ml anti-CD3 antibody (BD Pharmingen). After 3 days, Tresp cell proliferation was assessed by dilution of CTV fluorescence intensity with flow cytometry. For calibration of Tresp cell absolute numbers, CountBright Absolute Counting Beads (Thermo Fisher Scientific) were mixed with the cell samples before cell surface staining and assayed with flow cytometry. For the IFN-γ neutralization, 10 μg/ml anti-IFN-γ (eBioscience) or 10 μg/ml anti-IgG antibody (eBioscience) was added into culture medium.

### CD4^+^ T cell transfer model of colitis

CD4^+^CD25^−^CD45RB^hi^ naïve T cells (1 × 10^5^) from CD45.1^+^ C57BL/6 and CD45.2^+^CD4^+^FYP^+^ Treg cells (1 × 10^5^) from *Foxp3*^Cre/+^ or *Foxp3*^Cre/+^*IkE5*^f/f^ mice were mixed and intravenously transferred into *Rag2*^−/−^ recipients to induce colitis. Body weight of recipients was measured once a week up to day 42. Body weight change was assessed by the ratio of body weight to day 0 (Day 0 = 100%). At day 42, all mice were sacrificed and subjected to measurement of colon length, inspection of histology and flow cytometric analysis. For the IFN-γ neutralization, either anti-IFN-γ (250 μg/mouse) (BioXCell) or anti-IgG antibody (250 μg/mouse) (BioXCell) intraperitoneally injected into *Rag2*^−/−^ recipients every 3 days.

### Murine tumor models

*Foxp3*^eGFP-Cre-ERT2^ and *Foxp3*^eGFP-Cre-ERT2^*IkE5*^f/f^ mice were intradermally inoculated with B16F0 murine melanoma cells (5 × 10^5^) or MC38 murine colon adenocarcinoma cells (2 × 10^5^) into their shaved flanks (Day 0) and intraperitoneally injected with tamoxifen (4 mg/mouse in corn oil) (Sigma-Aldrich) on day 0, 1 and 3. Tumors were measured every 2 or 3 days with digital calipers and tumor area (mm^2^) was calculated as length x width. Tumor weight was measured on day 18. Tumors, draining lymph nodes (dLNs) and non-draining lymph nodes (ndLNs) were collected for analysis on day 10. Tumor-infiltrating leukocytes (TILs) were prepared with the Tumor Dissociation Kit (Miltenyi Biotec) and the gentleMACS Octo Dissociator (Miltenyi Biotec), according to the manufacturer’s instructions.

### RNA-seq and data analysis

Purified CD4^+^FYP^+^ Treg cells were stimulated with Cell Stimulation Cocktail (eBioscience) for 2 hrs and lysed in RLT buffer (Qiagen) containing 1% 2-mercaptoethanol (Thermo Fisher Scientific), followed by RNA reverse transcription by SMART-Seq v4 Ultra Low Input RNA Kit for Sequencing (Clontech). After enzymatic fragmentation of cDNA samples by KAPA Frag kit (KAPA Biosystems), sequencing libraries were prepared using KAPA Hyper Prep Kit (KAPA Biosystems). Sequencing of the cDNA libraries was performed by HiSeq2500 (Illumina).

Gene expression was quantified using ikra (v2.0), an RNA-seq pipeline centered on Salmon.^63^ The ikra pipeline executed the following tools for the quality control, trimming, and quantification of transcripts; Trim Galore (v0.6.6), Salmon (v1.4.0), and tximport (v1.6.0), consequently outputting the scalled TPM and TPM values. As the reference, GENCODE vM26 was used. The differentially expressed genes (DEGs) (Fold change, 2-fold; FDR < 0.1) were identified by DEseq2 in iDEP (v0.90)^64^ using the scalled TPM values. Gene set enrichment analysis (v4.0.3)^65^ was performed with the following settings: collapse = true, permutation type = gene_set, scoring = weighted. Hierarchical clustering was performed using the heatmap.2 function in R package gplots.

### ChIP-seq and data analysis

ChIP-seq experiments were performed as previously described with minor modification.^60^ Purified Treg cells were stimulated with Cell Stimulation Cocktail (eBioscience) for 1 hr at 37 ℃. Stimulated Treg cells were then cross-linked with 1% Formardehyde for 5 min (histone ChIP) or 30 min (transcription factor ChIP) at room temperature. After nuclear extraction, Cross-linked DNA was fragmentated by sonication using Digital Sonifier (Branson). The lysate was incubated overnight at 4 °C with 50-100 μl DynaBeads IgG magnetic beads (Thermo Fisher Scientific) that had been pre-incubated with 2.5-5 μg appropriate antibodies. Samples were washed, eluted, reverse cross-linked at 65 ℃ for 24 hrs, and purified using ChIP DNA Clean & Concentrator (Zymo Research). For the transcription factor ChIP-seq, purified ChIP DNA was fragmented using Covaris Focused-ultrasonicator S220 (Covaris) before library preparation. Library was prepared using Ion Xpress Plus Fragment Library Kit (Thermo Fisher Scientific) according to manufacturer’s instructions and sequenced by IonS5 sequencer system (Thermo Fisher Scientific). Antibodies used were anti-H3K27ac (GeneTex), anti-Foxp3 (Abcam), anti-Ikaros (Sigma-Aldrich), anti-p300 (Abcam) and anti-NFAT1 (Abcam).

For data analysis, the quality of sequence reads was confirmed using fastQC to confirm that the average of Phred score was over 20. Raw sequences were trimmed with fastx_trimmer in fastx_tool kit, using following setting; fastx_trimmer -Q33 -f 12 −l 220 -i ${name}.fastq | fastq_quality_trimmer -Q33 -t 20 −l 30 -o ${name}_trimm.fastq. Sequencing reads were mapped to the mouse genome mm10 using Bowtie2 with default setting; bowtie2 -p 30 -x mm10 -U ${name}_trim.fastq -S ${name}_accept.sam. For visualization of ChIP peaks, peak call was performed using MACS2^66^ by the following command; macs2 callpeak -t ${name}_accept.sam -c input.sam -g mm -n ${name} -B -q 0.01 –nomodel. Bdg2bw tool used for the conversion of bedgraph to bigwig. Integrated genome viewer (IGV) was used for the visualization of peak or region data using group-auto scaling, auto-scaling based on normalized to total mapped reads. ChIP-seq peaks were defined using FindPeaks, with their size fixed at 500 base pairs, minimum distance between peaks being 500 base pairs, FDR set as default, and local filtering switched off. The overlap of among ChIP-seq peaks was defined by ≥ 1 base pair overlap using Bedtools. To identify common or specific peaks between given conditions, the identified peaks in each condition were merged. Then, the merged peaks were used to define common or specific peaks using HOMER (getDifferentialPeaks) with default parameters. Normalized density plots of histone modifications peaks as well as binding of transcription factors around Foxp3-binding sites were calculated using annotatePeaks in Homer package.^67^

### ATAC-seq and data analysis

ATAC-seq was performed as previously described.^61^ Briefly, purified CD4^+^FYP^+^ Treg cells (1 × 10^5^) were lysed using 50 μl of lysis buffer (0.01% digitonin, 0.1% NP-40, 0.1% Tween 20 in resuspension buffer; 10 mM Tris-HCl pH7.5, 100 mM NaCl, 3 mM MgCl2) for 3 min on ice. After removing lysis buffer by centrifugation, Tn5 tagmentation was performed using Illumina Tagment DNA TDE1 Enzyme and Buffer Kits (Illumina) at 37℃ for 30 min, with shaking at 1000 rpm following manufacturer’s instruction. After purification using DNA Clean & Concentrator-5 (Zymo Research), tagmented DNA was amplified using NEBNext High-Fidelity PCR Master Mix (New England BioLabs) with the following primers: 5’-CAAGCAGAAGACGGCATACGAGATNNNNNNNNGTCTCGTGGGCTCGGAGATG T-3’ and 5’-AATGATACGGCGACCACCGAGATCTACACNNNNNNNNTCGTCGGCAGCGTCAG ATGTG-3’ (barcode sequences are indicated as NNNNNNNN). Prepared DNA libraries were size-selected (150-1000 bp) by Ampure XP (Beckman Coulter). Sequencing was performed using NextSeq500 (Illumina).

For ATAC-seq analysis, sequenced reads were processed with fastx_trimmer and cmpfastq_pe. Processed reads were mapped to mm10 reference using Bowtie2 with default setting. The peak call was performed using MACS2^66^. ATAC-seq peaks were defined using FindPeaks, with their size fixed at 250 base pairs, FDR set as default, and local filtering switched off. Normalized density of mapped reads around Foxp3-binding sites was determined using annotatePeaks.^67^

### Retroviral transduction

Purified CD4^+^FYP^+^ Treg cells (1 × 10^5^) were stimulated the plate-bound α-CD3 (1 μg/ml) (BD Pharmingen), α-CD28 (1 μg/ml) (BD Pharmingen) and IL-2 (100 U/ml) for 24 hrs. After activation, fresh retrovirus supernatant was added and the cells were spun with 2500 rpm for 90 min at 32°C. After spin infection, the cells were cultured under the above conditions and harvested on day 4 for flow cytometric analysis.

### *In vitro* CRISPR/Cas9-mediated gene targeting

CRISPR RNAs (crRNAs) for target genes were designed using the Integrated DNA Technologies (IDT) guide RNA design tool. Two or three crRNAs per target were designed and mixed to use. Negative control crRNA and tracrRNA were purchased from IDT. Purified CD4^+^FYP^+^ Treg cells (1 × 10^5^) were cultured with Dynabeads mouse CD3/CD28 T cell stimulator (25 μl/ml) (Thermo Fisher Scientific) and IL-2 (1500 U/ml) for 7 days. Expanded Treg cells (2 × 10^6^) were resuspended into P4 Primary Cell solution (Lonza Bioscience) and nucleofected with Cas9 protein and gRNAs, which are mixture of crRNAs and tracrRNA, at the DG137 program by Amaxa 4D (Lonza Bioscience). After nucleofection, cells were washed and re-cultured with Dynabeads mouse CD3/CD28 T cell stimulator (25 μl/ml) (Thermo Fisher Scientific) and IL-2 (1500 U/ml) for 3 days, followed by staining for surface and intracellular molecules.

### Data and code availability

All data generated in this study are included in this published article and online supplemental materials. The RNA-seq, the ChIP-seq and the ATAC-seq data have been deposited in the GEO database^68^ under the accession codes: GSE229592. Any additional information required to reanalyze the data reported in this paper is available from the lead contact upon request.

### Quantification and statistical analysis

Statistical analyses were calculated with Prism software (GraphPad). Normal distribution was assumed a priori for all samples. *P* values of less than 0.05 were considered significant (\**P* < 0.05; \*\**P* < 0.01; \*\*\**P* < 0.001).

## ACKNOWLEDGMENTS

We thank Fujii Memorial Research Institute for providing REF001329, J. White for reading the manuscript, A. Shimonishi, M. Zaizen and R. Ishii for help with mice and cell sorting. Bioinformatics analyses were conducted using the computer system at the Genome Information Research Center of the Research Institute for Microbial Diseases at Osaka University. This study was funded in part by grants-in-aid from the Ministry of Education, Sports, and Culture of Japan (16H06295), and Japan Agency for Medical Research and Development (P-CREATE; 18cm0106303, and LEAP; 18gm0010005) to S.S. and the Japan Society for the Promotion of Science (JSPS) KAKENHI Grant-in-Aid for Scientific Research (B) 21H02748, and the Takeda Science Foundation to K.I.

## AUTHOR CONTRIBUTIONS

K.I. and S.S. designed the research and analyzed the data. K.I. performed most of experiments and bioinformatic analysis of next generation sequencing. Y.N. supported to run next generation sequencing. Y.H. and T.K. performed most of human experiments. L.J., Y.K. and C.K. participated in specific experiments. K.I. prepared the manuscript. K.G. and S.S. critically revised the manuscript.

---

**Figure S1.**
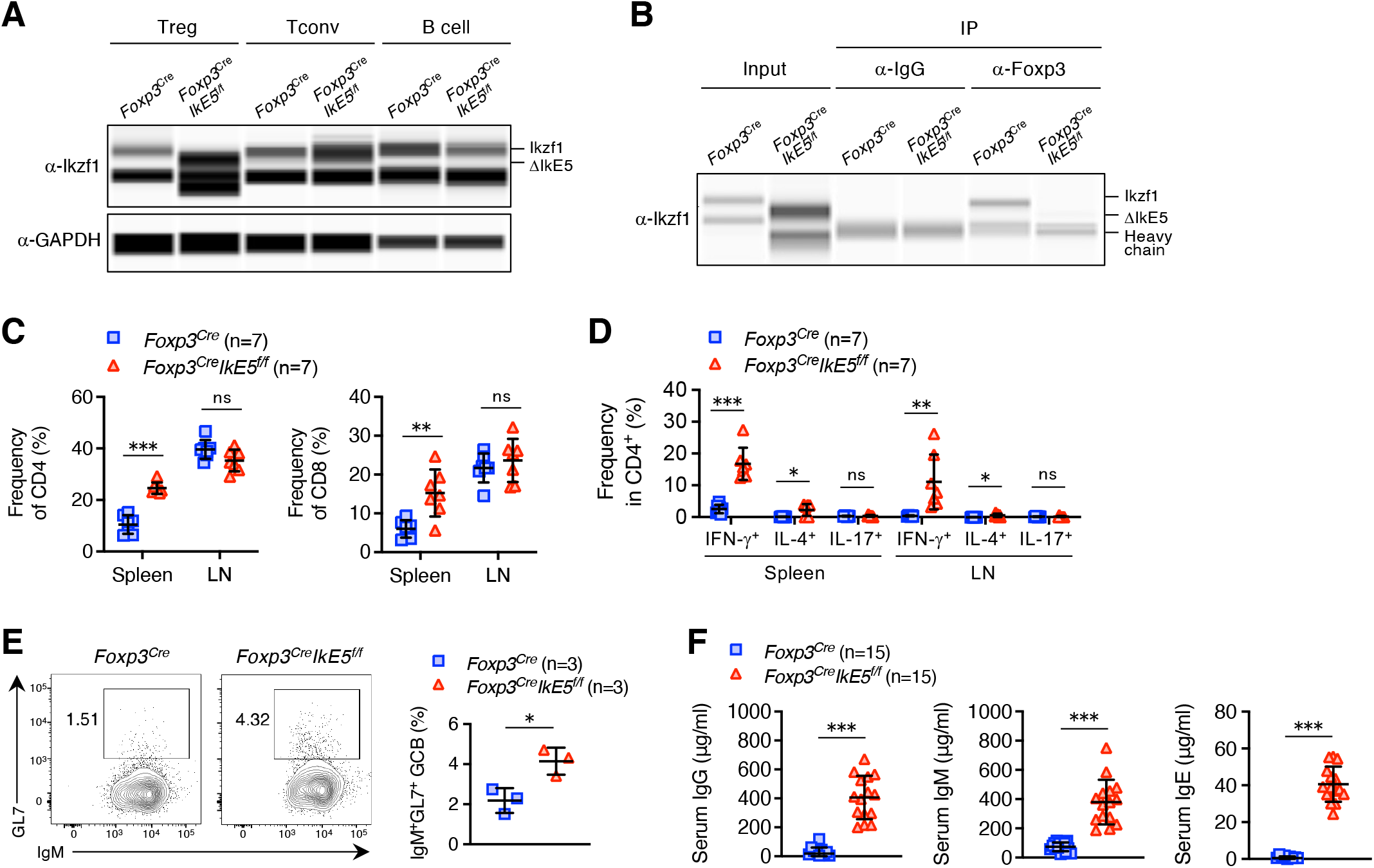
Treg-specific *IkE5*-deficient mice show immune activation. (A) Immunoblot analysis of Ikzf1 and GAPDH in Treg, Tconv and B cells from *Foxp3*^Cre^ and *Foxp3*^Cre^*IkE5*^f/f^ mice. The shift in band size indicates *IkE5*-deletion. (B) A representative image of interaction between Foxp3 and Ikzf1 in CD4^+^YFP^+^ Treg cells from *Foxp3*^Cre^ and *Foxp3*^Cre^*IkE5*^f/f^ mice by Co-IP. (C) Frequencies of CD4^+^ and CD8^+^ T cells in the spleen and LN from *Foxp3*^Cre^ and *Foxp3*^Cre^*IkE5*^f/f^ mice at 3 to 4 weeks of age (mean ± SD, *n* = 7 per group). (D) Frequencies of indicated cytokine-producing CD4^+^ T cells in the spleen and LN from *Foxp3*^Cre^ and *Foxp3*^Cre^*IkE5*^f/f^ mice at 3 to 4 weeks of age (mean ± SD, *n* = 7 per group). (E) Frequencies of B220^+^IgM^+^GL7^+^ GCB cells in the spleen from *Foxp3*^Cre^ and *Foxp3*^Cre^*IkE5*^f/f^ mice at 3 to 4 weeks of age (right)(mean ± SD, *n* = 3 per group). A representative flow cytometry plot of B220^+^ splenocytes in the above mice (left). (F) Concentration of anti-IgG, anti-IgM and anti-IgE antibodies in serum of *Foxp3*^Cre^ and *Foxp3*^Cre^*IkE5*^f/f^ mice at 3 to 4 weeks of age, determined by ELISA (mean ± SD, *n* = 15 per group). Data are representative of at least three independent experiments (A,B) or summary of at least three independent experiments (C-F). *P* values determined by unpaired *t*-tests followed by Holm-Šídák multiple comparisons test (C,D) or two-tailed unpaired *t*-tests (E,F). ns, not significant, \**P*< 0.05; \*\**P* < 0.01; \*\*\**P* < 0.001.

**Figure S2.**
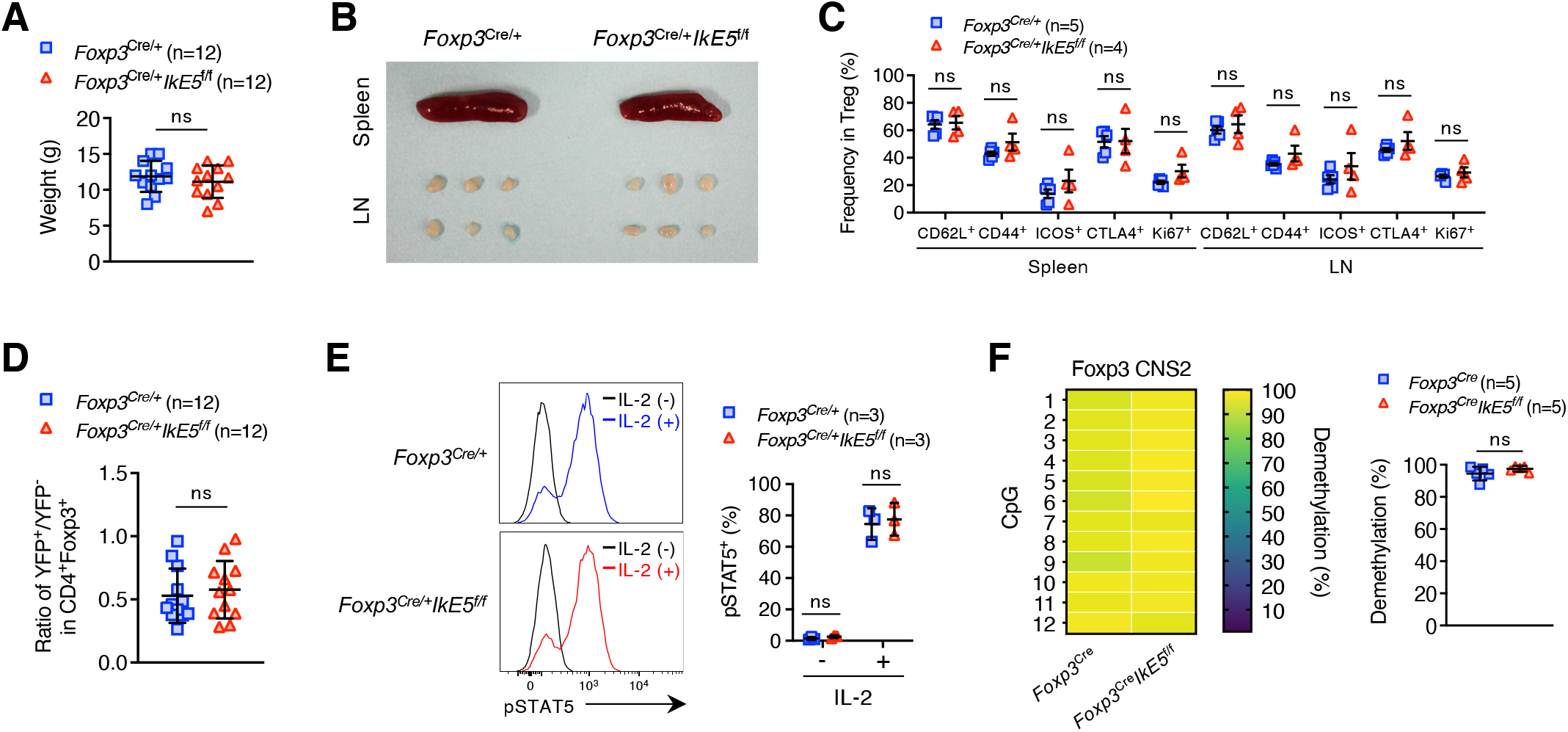
*IkE5*-deletion does not affect the activation status, IL-2/STAT5 signaling and DNA demethylation of Treg cells. (A) Body weight of *Foxp3*^Cre/+^ and *Foxp3*^Cre/+^*IkE5*^f/f^ mice at 3 to 4 weeks of age (mean ± SD, *n* = 12 per group). (B) A representative appearance of spleen and peripheral lymph nodes (LN) from *Foxp3*^Cre/+^ and *Foxp3*^Cre/+^*IkE5*^f/f^ mice at 3 to 4 weeks of age. (C) Frequencies of indicated molecule-expressing Treg cells in the spleen and LN from *Foxp3*^Cre/+^ (*n* = 5) and *Foxp3*^Cre/+^*IkE5*^f/f^ (*n* = 4) mice at 3 to 4 weeks of age (mean ± SD). (D) The ratio of YFP^+^/YFP^−^ Treg cells in the thymus from *Foxp3*^Cre/+^ (*n* = 12) and *Foxp3*^Cre/+^*IkE5*^f/f^ (*n* = 12) mice at 3 to 4 weeks of age (mean ± SD). (E) Purified CD4^+^YFP^+^ Treg cells from *Foxp3*^Cre/+^ and *Foxp3*^Cre/+^*IkE5*^f/f^ mice at 3 to 4 weeks of age were first starved for 45 min at 37°C, followed by stimulation with IL-2 (100 U/ml) for 30 min at 37°C. A representative flow cytometry histogram of pSTAT5 (left) and frequencies of pSTAT5^+^ population (right) in stimulated Treg cells (mean ± SD, *n* = 3 per group). (F) CpG methylation status within *Foxp3* CNS2 region in CD4^+^YFP^+^ Treg cells from *Foxp3*^Cre^ and *Foxp3*^Cre^*IkE5*^f/f^ mice at 3 to 4 weeks of age (mean ± SD, *n* = 5 per group). Data are summary of at least three independent experiments (A,C-F) or representative of three independent experiments (B). *P* values determined by unpaired *t*-tests followed by Holm-Šídák multiple comparisons test (C,E) or two-tailed unpaired *t*-tests (A,D,F). ns, not significant.

**Figure S3.**
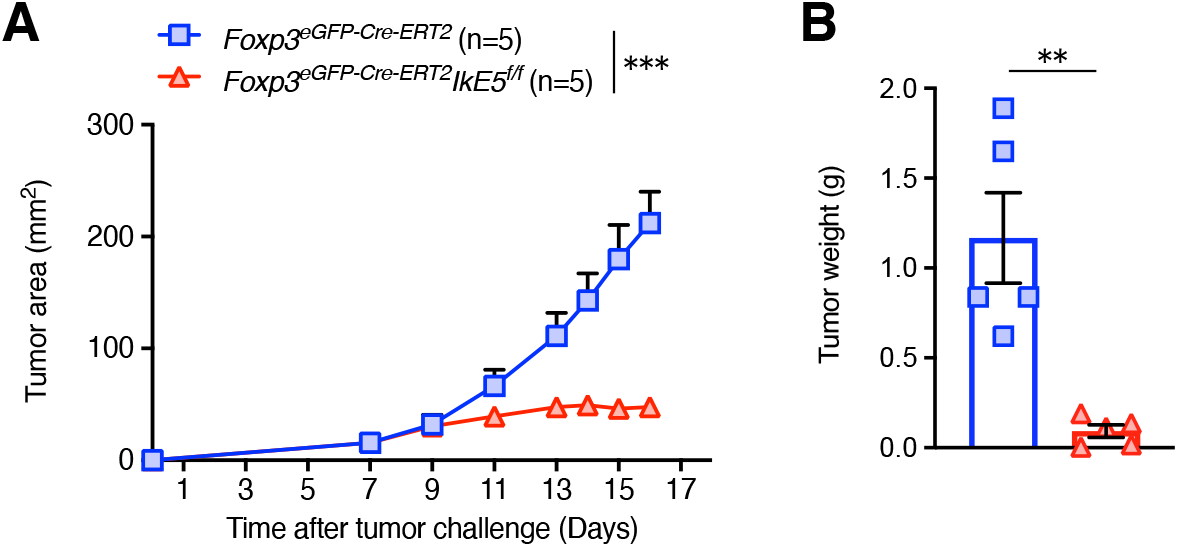
Treg-specific *IkE5*-deficient mice suppress progression of B16 melanoma. (A) *Foxp3*^eGFP-Cre-ERT2^ and *Foxp3*^eGFP-Cre-ERT2^*IkE5*^f/f^ mice were intradermally inoculated with B16F0 murine melanoma cells (5 × 10^5^) into their shaved flanks (Day 0) and intraperitoneally injected with tamoxifen on day 0, 1 and 3. Tumor area (mm^2^) was measured over 16 days (mean ± SEM, *n* = 5 per group). (B) Tumor weight in *Foxp3*^eGFP-Cre-ERT2^ and *Foxp3*^eGFP-Cre-ERT2^*IkE5*^f/f^ mice inoculated as indicated in (A) was measured on 16 days (mean ± SEM, *n* = 5 per group). Data are representative of two independent experiments (A,B). *P* values determined by ordinary two-way ANOVA followed by Šídák multiple comparisons test (A) or two-tailed unpaired *t*-tests (B). \*\**P*< 0.01; \*\*\**P* < 0.001.

**Figure S4.**
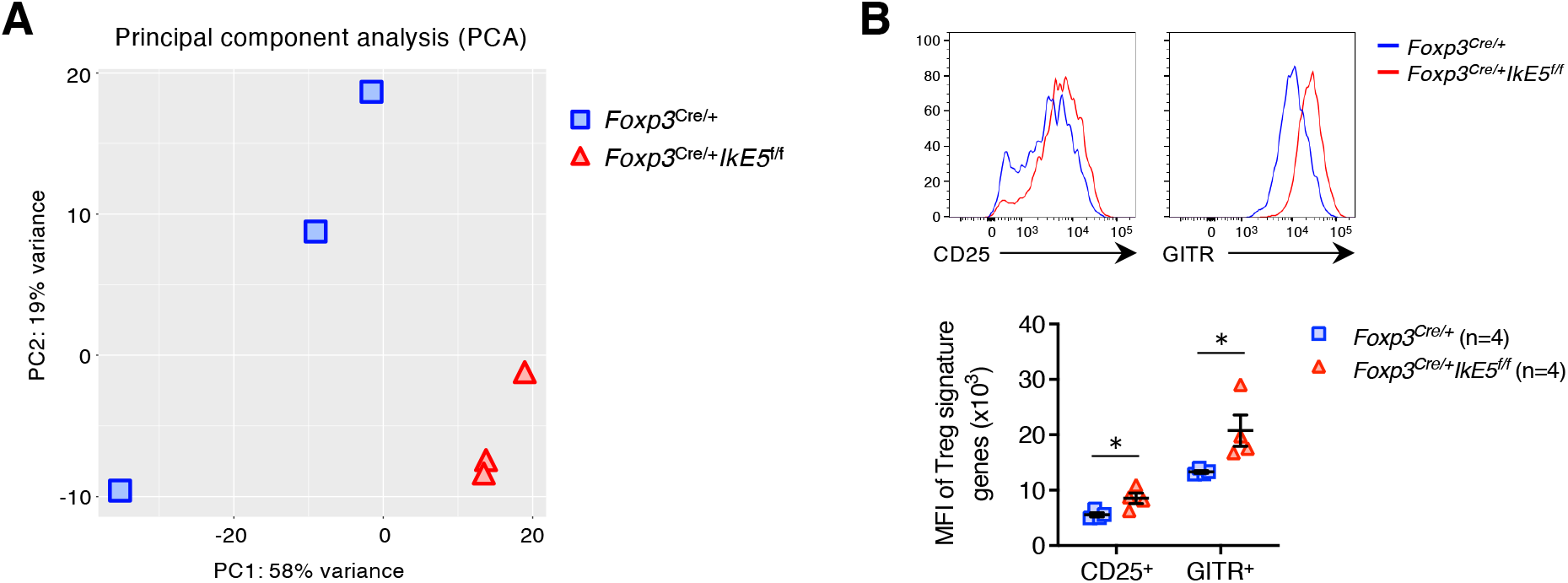
RNA-seq analysis in CD4^+^YFP^+^ Treg cells from *Foxp3*^Cre/+^ and *Foxp3*^Cre/+^*IkE5*^f/f^ mice. (A) Principal component analysis (PCA) of gene expression in CD4^+^YFP^+^ Treg cells from *Foxp3*^Cre/+^ and *Foxp3*^Cre/+^*IkE5*^f/f^ mice at 3 to 4 weeks of age (*n* = 3 per group). (B) Representative flow cytometry histograms (up) and the mean fluorescence intensity (MFI)(bottom) of indicated Treg signatures in CD4^+^YFP^+^ Treg cells from *Foxp3*^Cre/+^ and *Foxp3*^Cre/+^*IkE5*^f/f^ mice at 3 to 4 weeks of age (mean ± SEM, *n* = 4 per group). Data are representative of two independent experiments (A) or summary of three independent experiments (B). *P* values determined by unpaired *t*-tests followed by Holm-Šídák multiple comparisons test (B). \**P*< 0.05.

**Figure S5.**
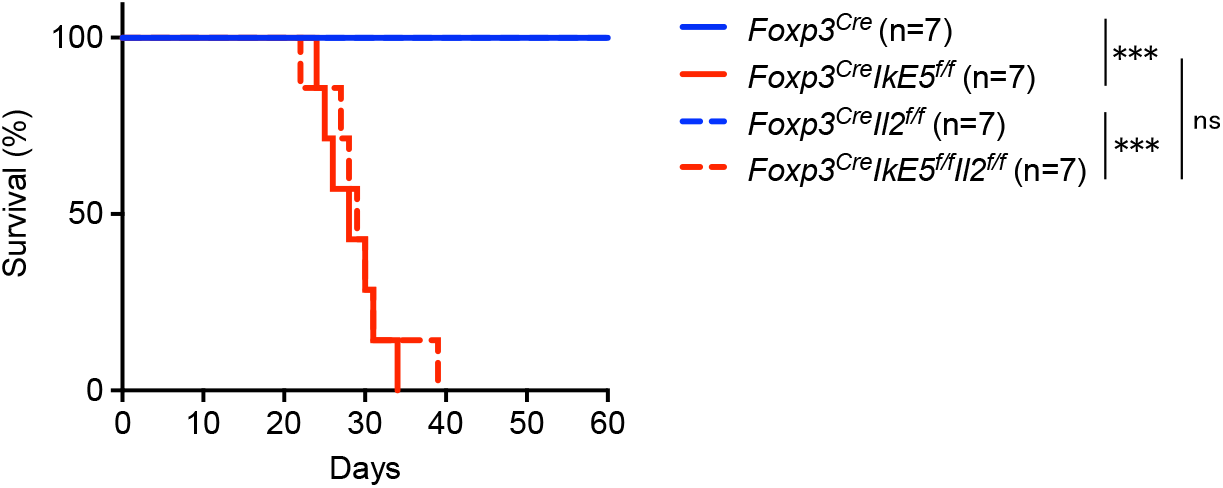
IL-2 is not responsible for the fatal systemic autoimmunity in *Foxp3*^Cre^*IkE5*^f/f^ mice. Kaplan-Meier survival curve of *Foxp3*^Cre^, *Foxp3*^Cre^*IkE5*^f/f^, *Foxp3*^Cre^*Il2*^f/f^ and *Foxp3*^Cre^*IkE5*^f/f^*Il2*^f/f^ mice (*n* = 7 per group). Data are summary of two independent experiments. *P* values determined by log-rank test. ns, not significant, \*\*\**P* < 0.001.

**Figure S6.**
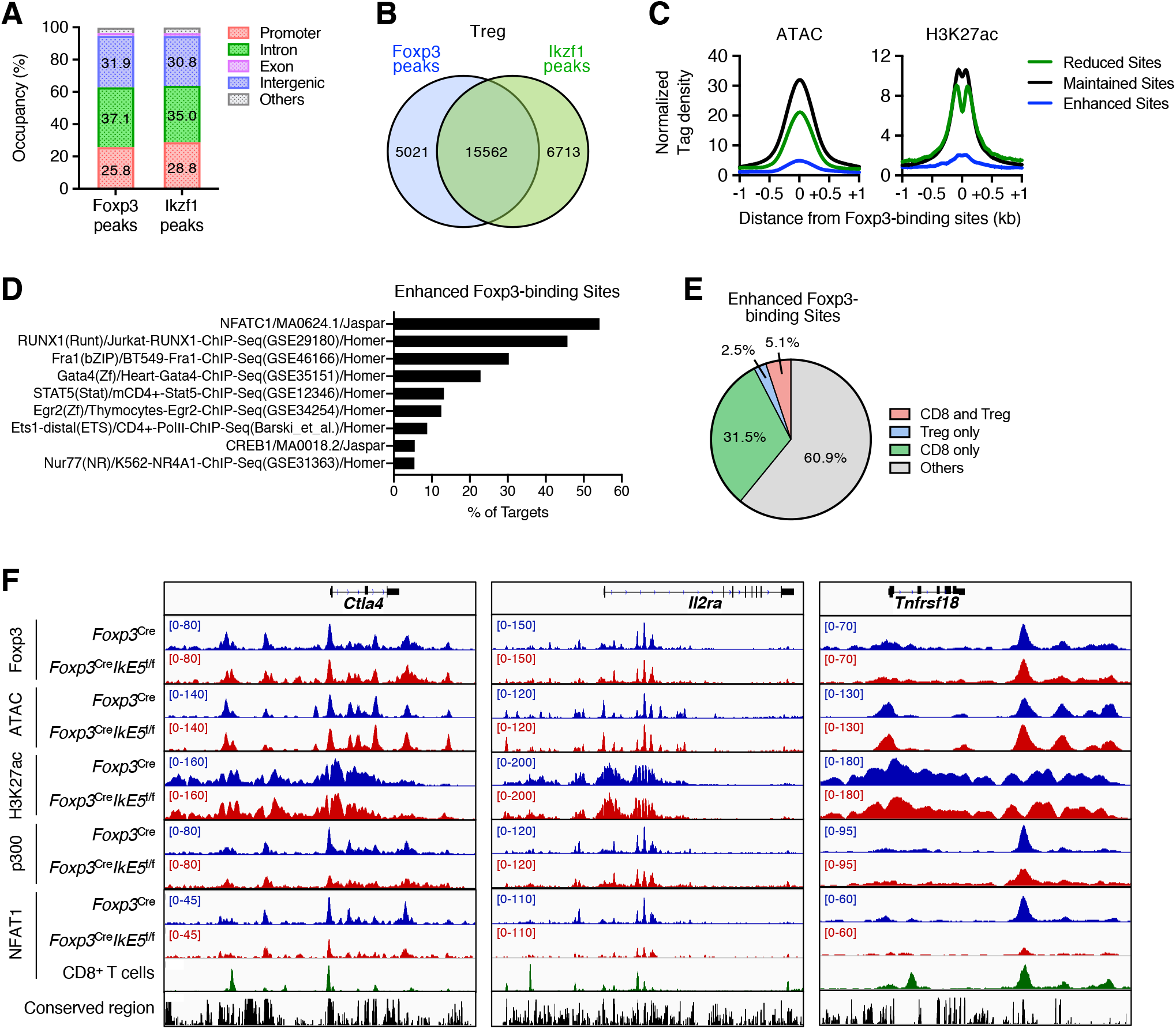
ChIP-seq analysis in CD4^+^YFP^+^ Treg cells from *Foxp3*^Cre^ and *Foxp3*^Cre^*IkE5*^f/f^ mice. (A) Peak annotation of Foxp3 and Ikzf1 ChIP-seq peaks in CD4^+^YFP^+^ Treg cells from *Foxp3*^Cre^ mice. (B) Venn diagram of Foxp3 (blue) and Ikzf1 (green) ChIP-seq peaks in CD4^+^YFP^+^ Treg cells from *Foxp3*^Cre^ mice. (C) Normalized density plots of ATAC and H3K27ac peaks around the Reduced Foxp3-binding Sites (green), Maintained Foxp3-binding Sites (black) and Enhanced Foxp3-binding Sites (blue) in CD4^+^YFP^+^ Treg cells from *Foxp3*^Cre^ mice. Normalized signal density is plotted within a window ± 1 kb centered on Foxp3-binding sites. (D) Motif enrichment analysis on the Enhanced Foxp3-binding Sites. (E) Pie chart illustrated the percentage of NFAT1-binding within the Enhanced Foxp3-binding Sites in CD8^+^ T and Treg cells. (F) Foxp3, p300, NFAT1, H3K27ac ChIP-seq and ATAC-seq signal tracks at the Treg up-regulated genes, such as *Ctla4*, *Il2ra* and *Tnfrsf18* genes loci in CD4^+^YFP^+^ Treg cells from *Foxp3*^Cre^ (blue) and *Foxp3*^Cre^*IkE5*^f/f^ (red) mice. Data of CD8^+^ T cells (green) is from the previous report.^41^ Sequence conservation among vertebrates (black) is also shown. Data are representative of two independent experiments (A-F).

**Figure S7.**
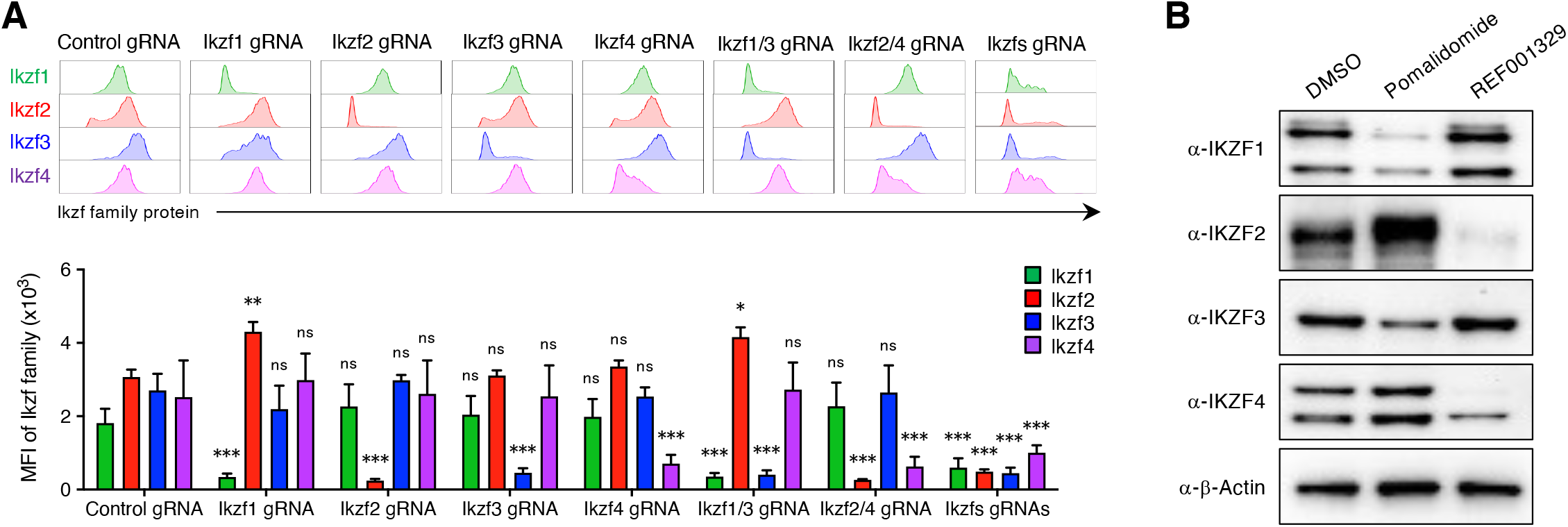
Confirmation of Ikzf family proteins disruption by the *in vitro* CRISPR/Cas9 system and thalidomide analogues. (A) A representative flow cytometry histogram (up) and MFI (bottom) of Ikzf family proteins in CD4^+^Foxp3^+^ Treg cells in Figure 7A (mean ± SD, *n* = 3 per group). (B) Immunoblot analysis of IKZF family proteins in human CD4^+^CD25^+^CD45RA^+^ and CD4^+^CD25^hi^CD45RA^−^ Treg cells stimulated with DMSO, Pomalidomide (10 μM) or REF001329 (10 μM) for 9 days. Data are summary of three independent experiments (A) or representative of three independent experiments (B). *P* values determined by two-way ANOVA followed by Dunnett’s multiple comparisons test (A). ns, not significant, \**P*< 0.05; \*\**P* < 0.01; \*\*\**P* < 0.001.

**Table S1.**
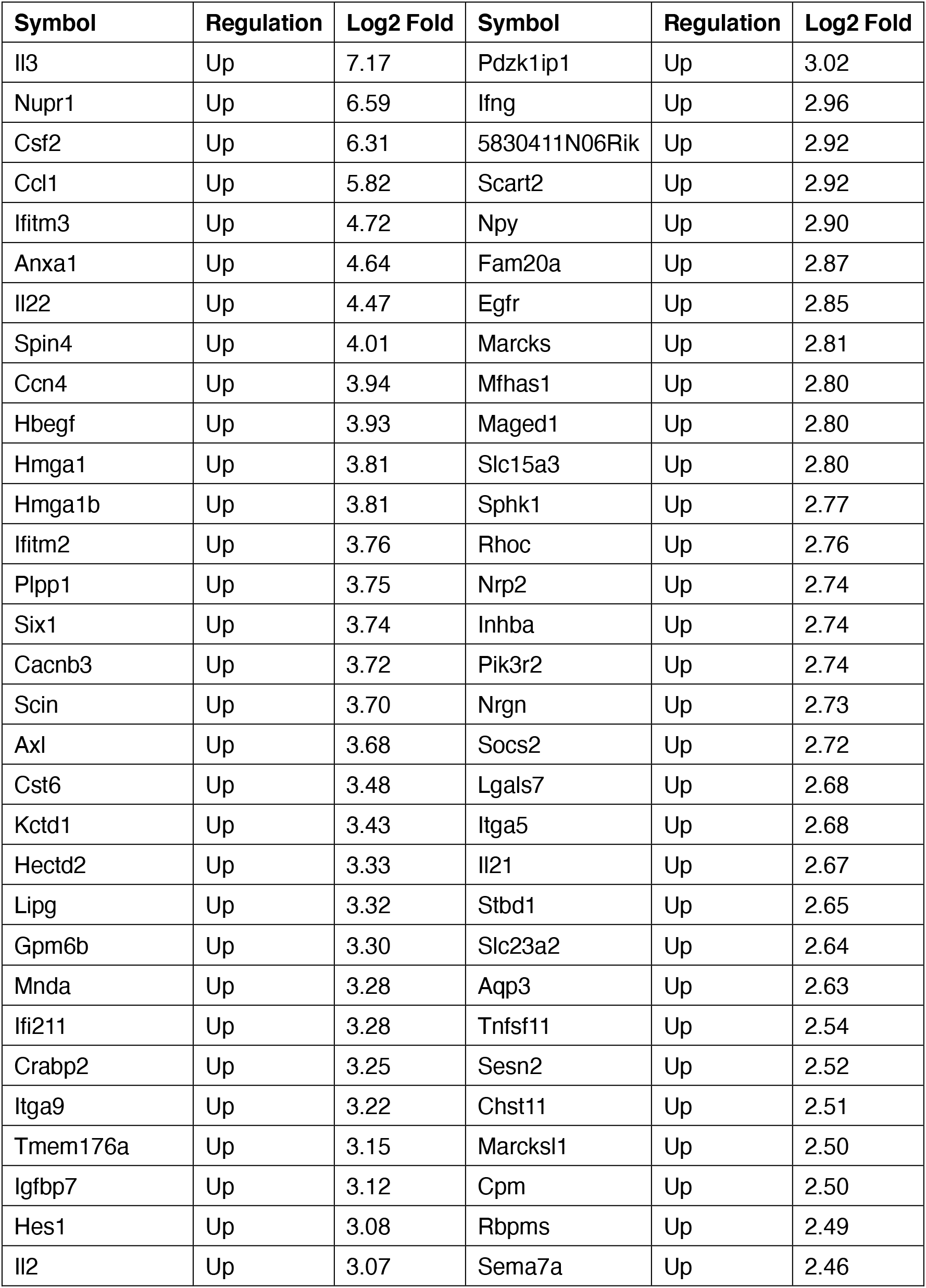
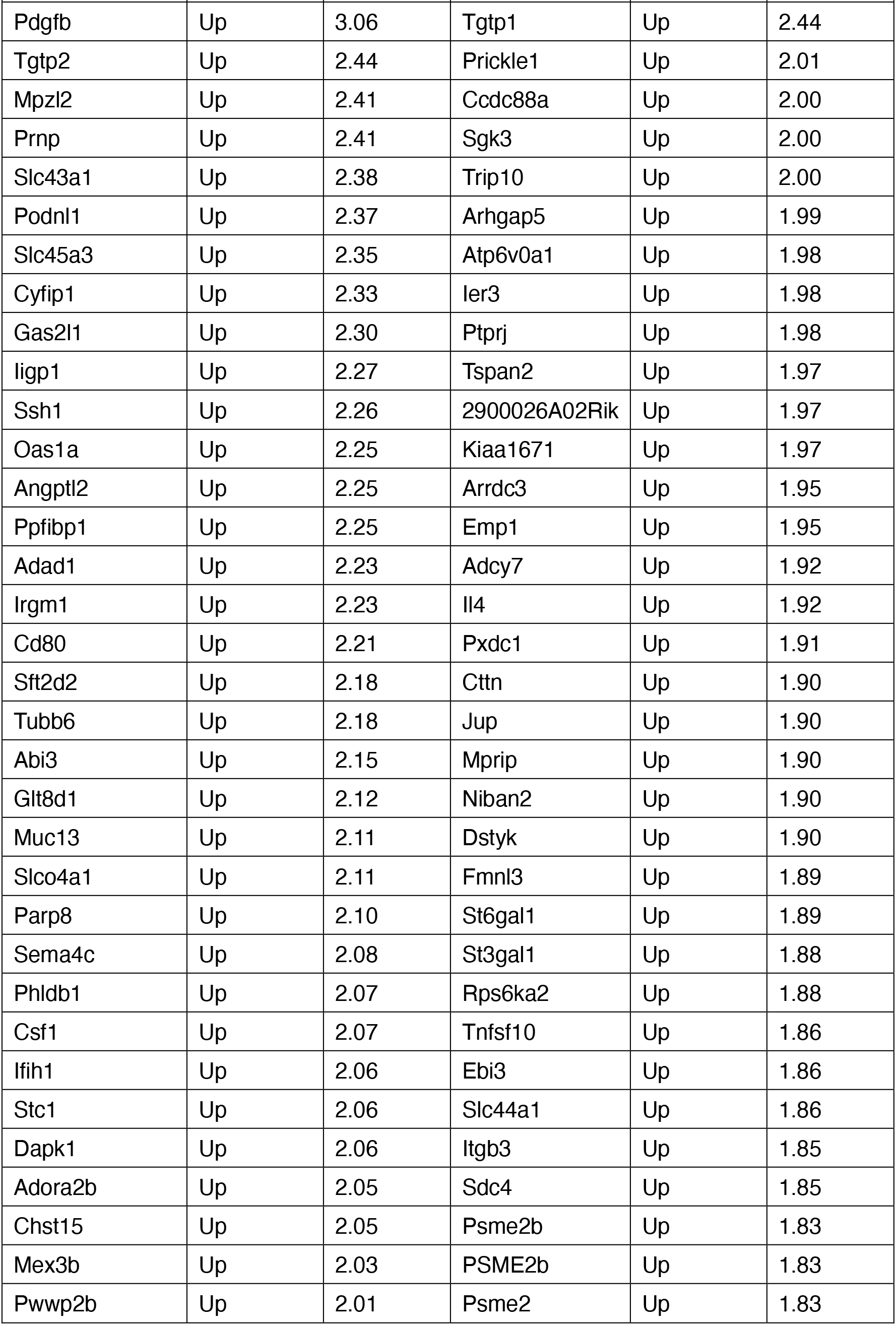
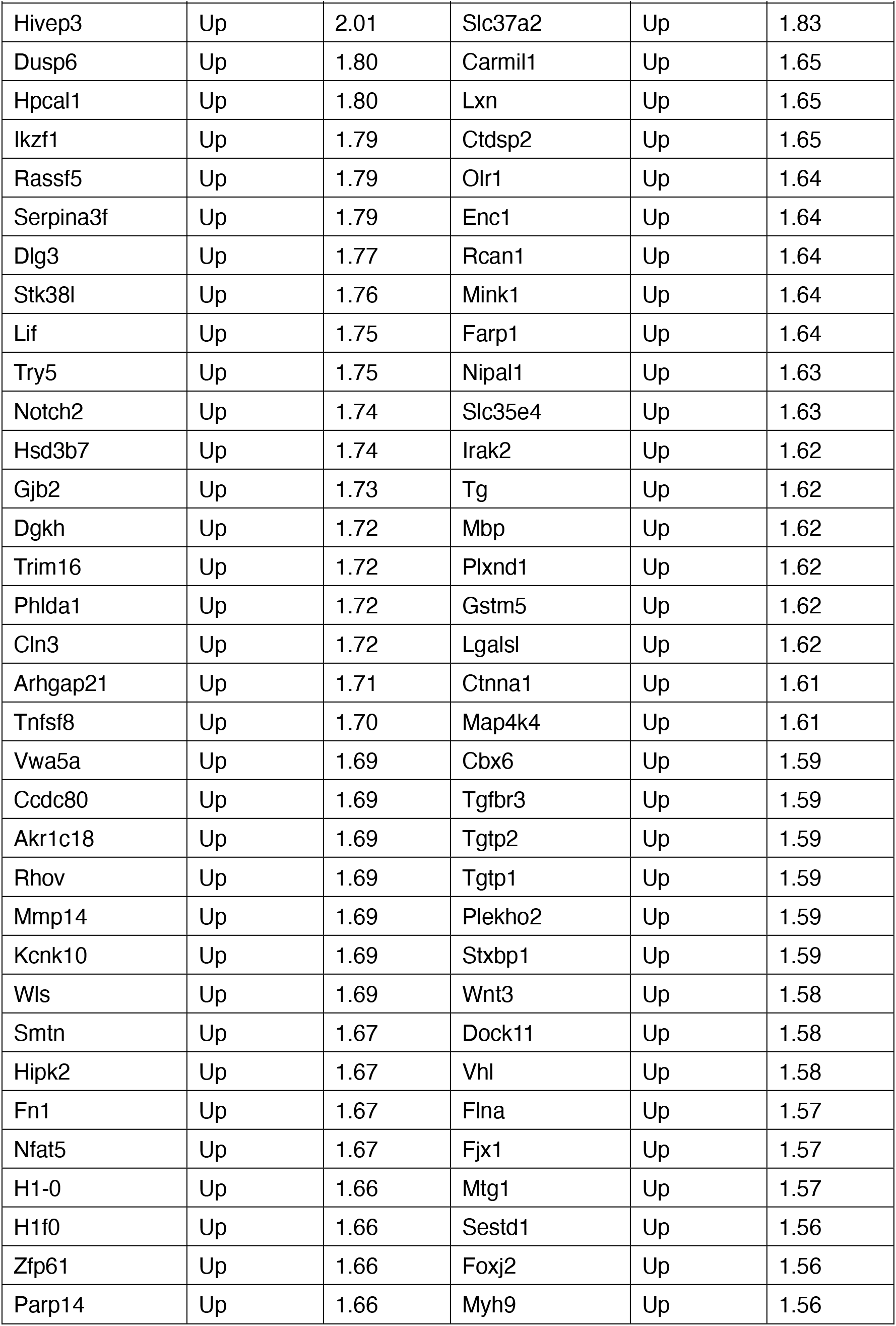
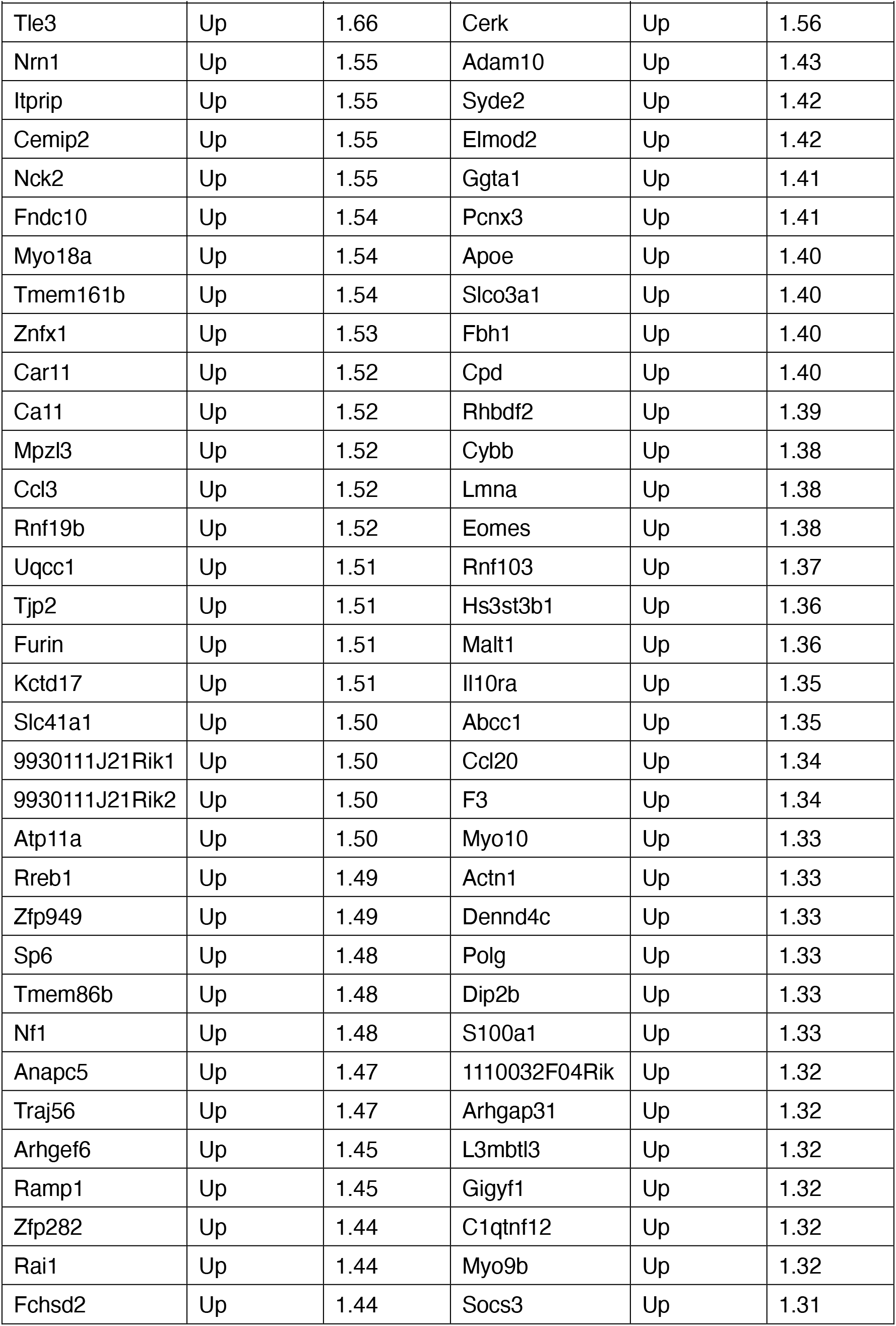
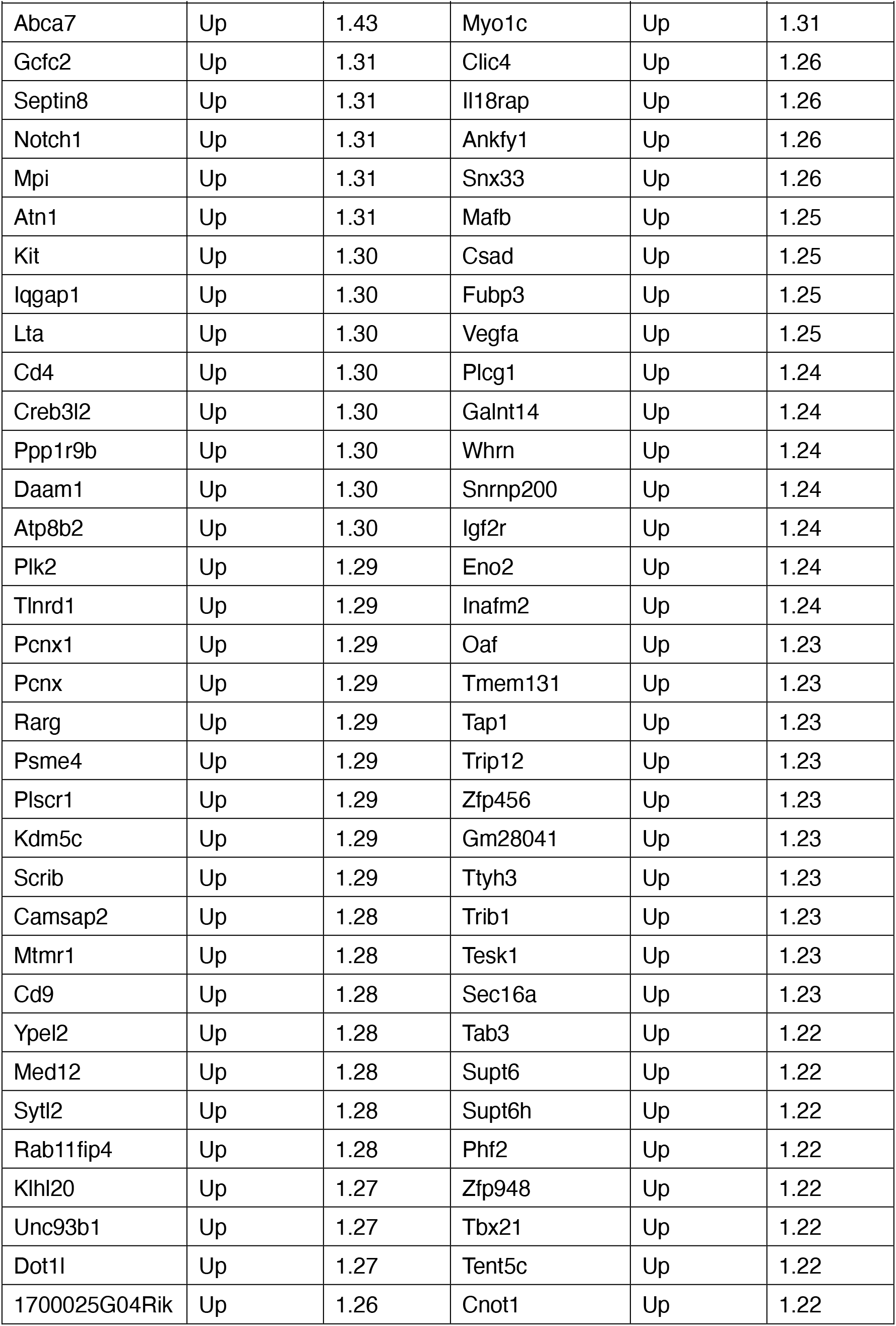
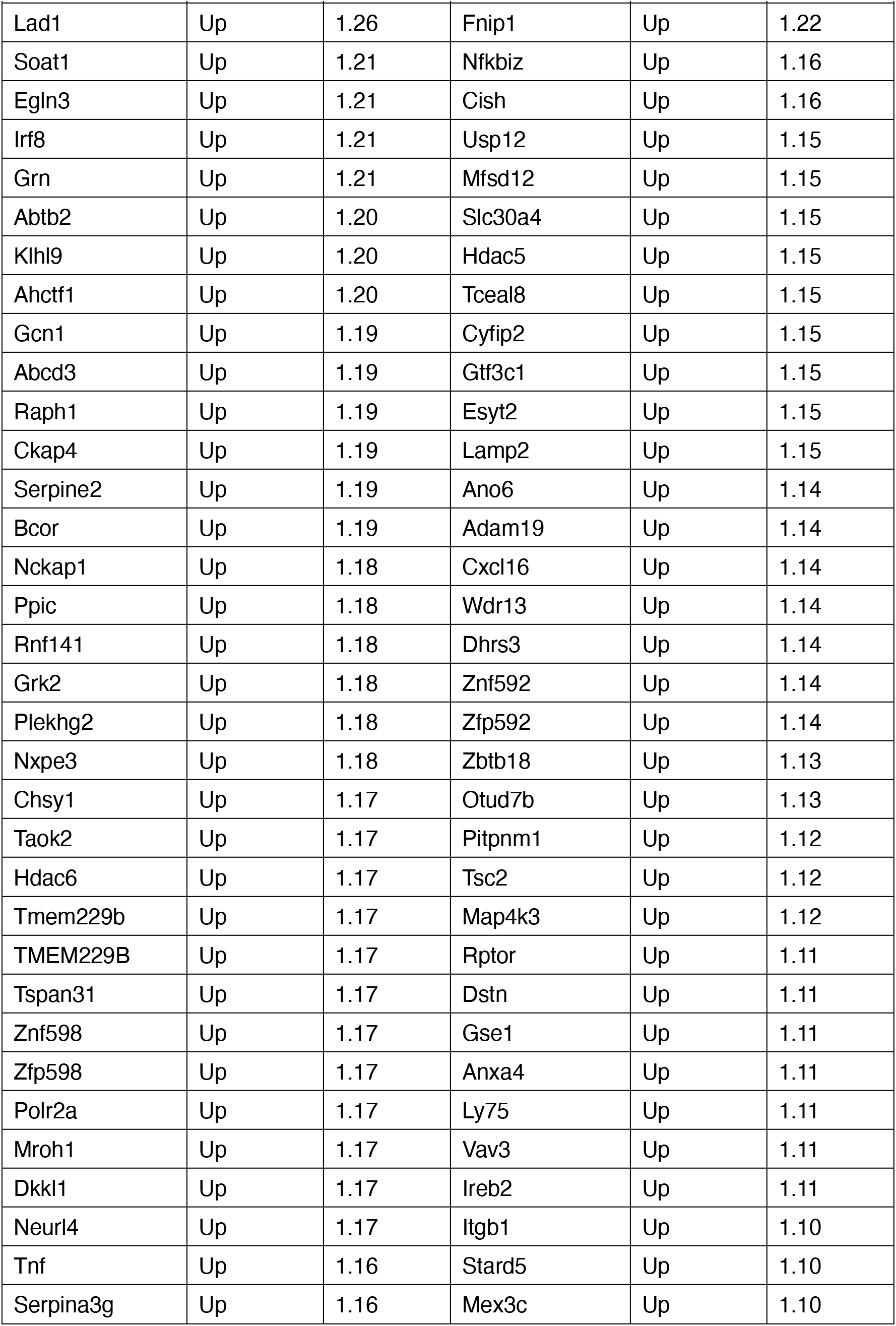
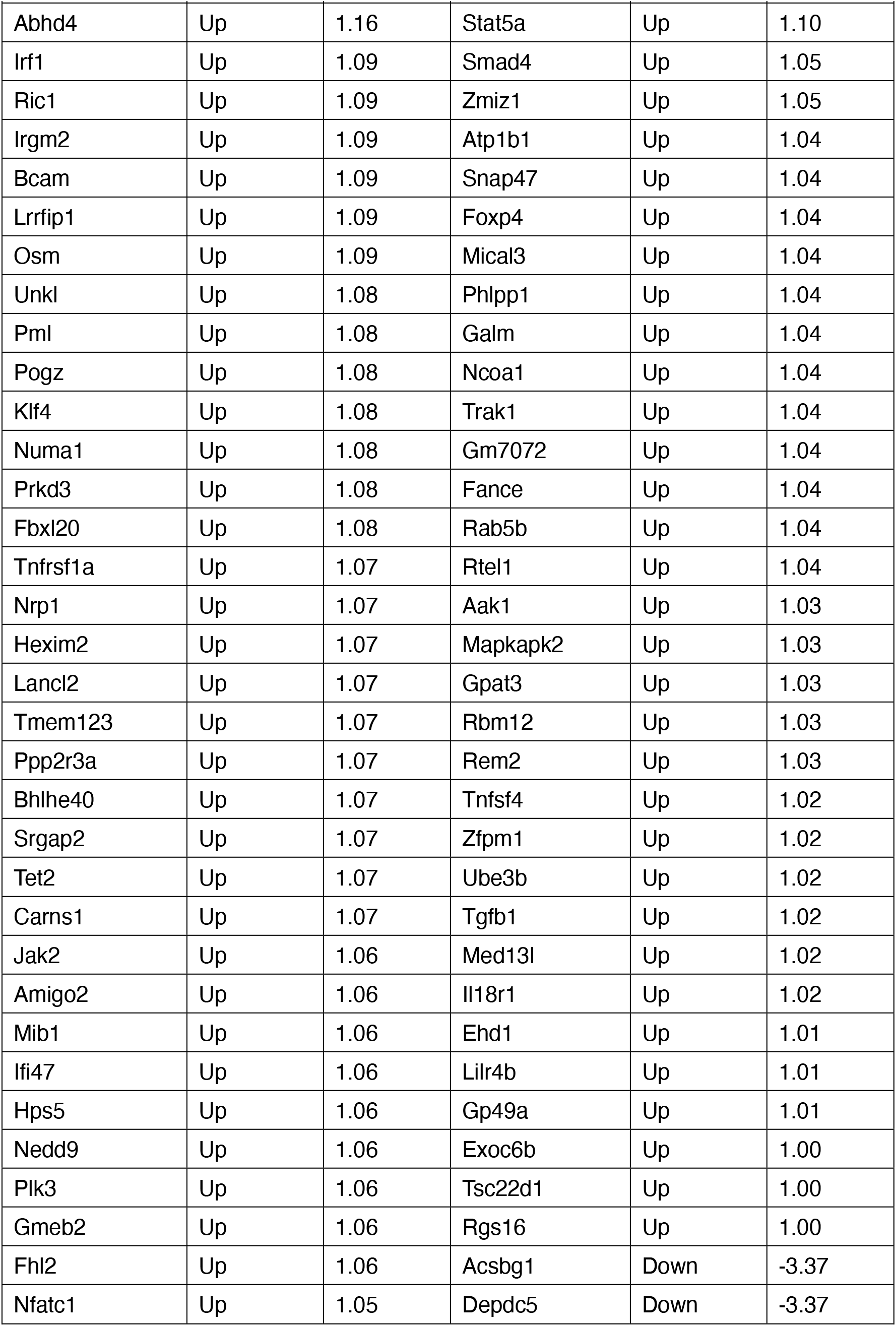
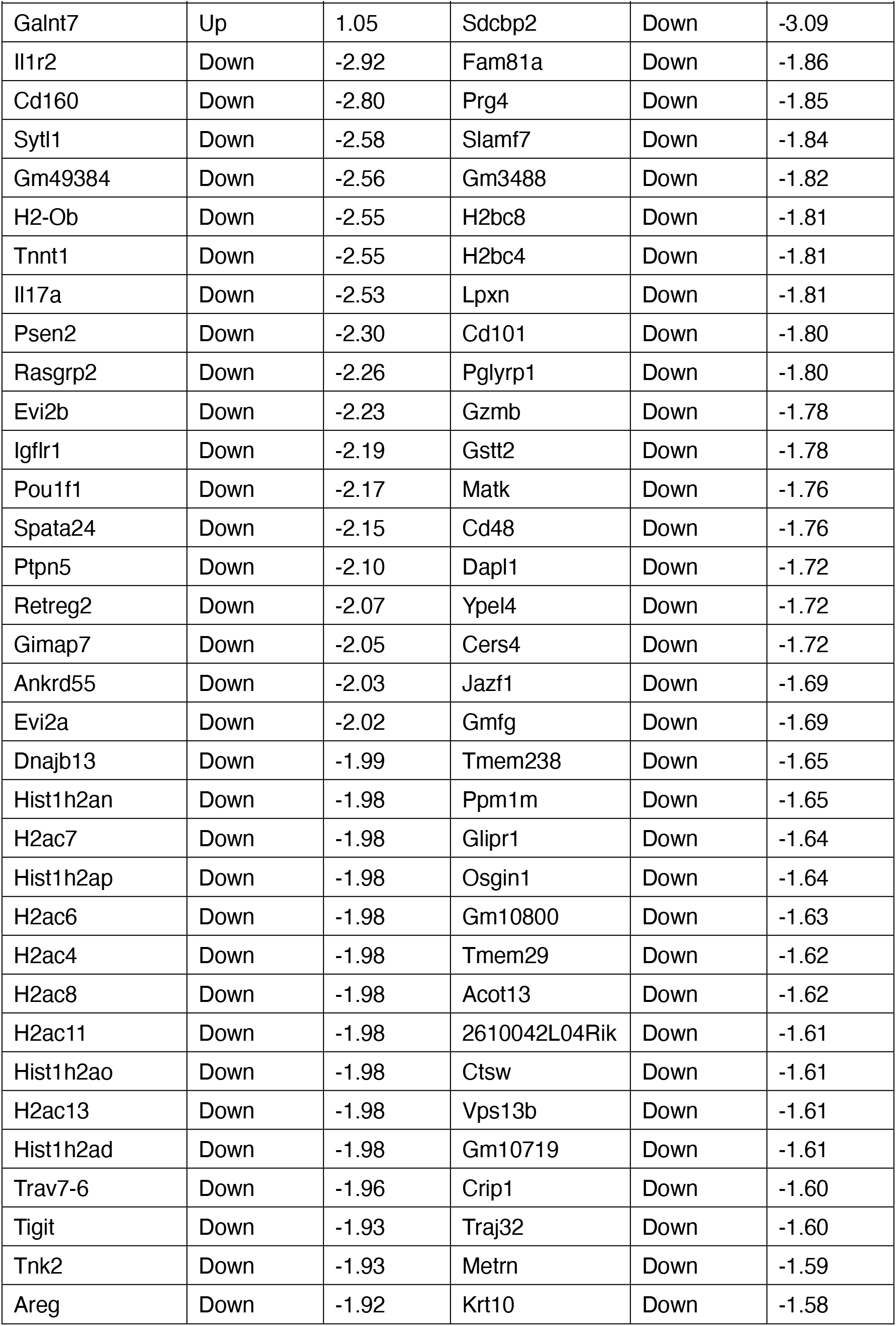
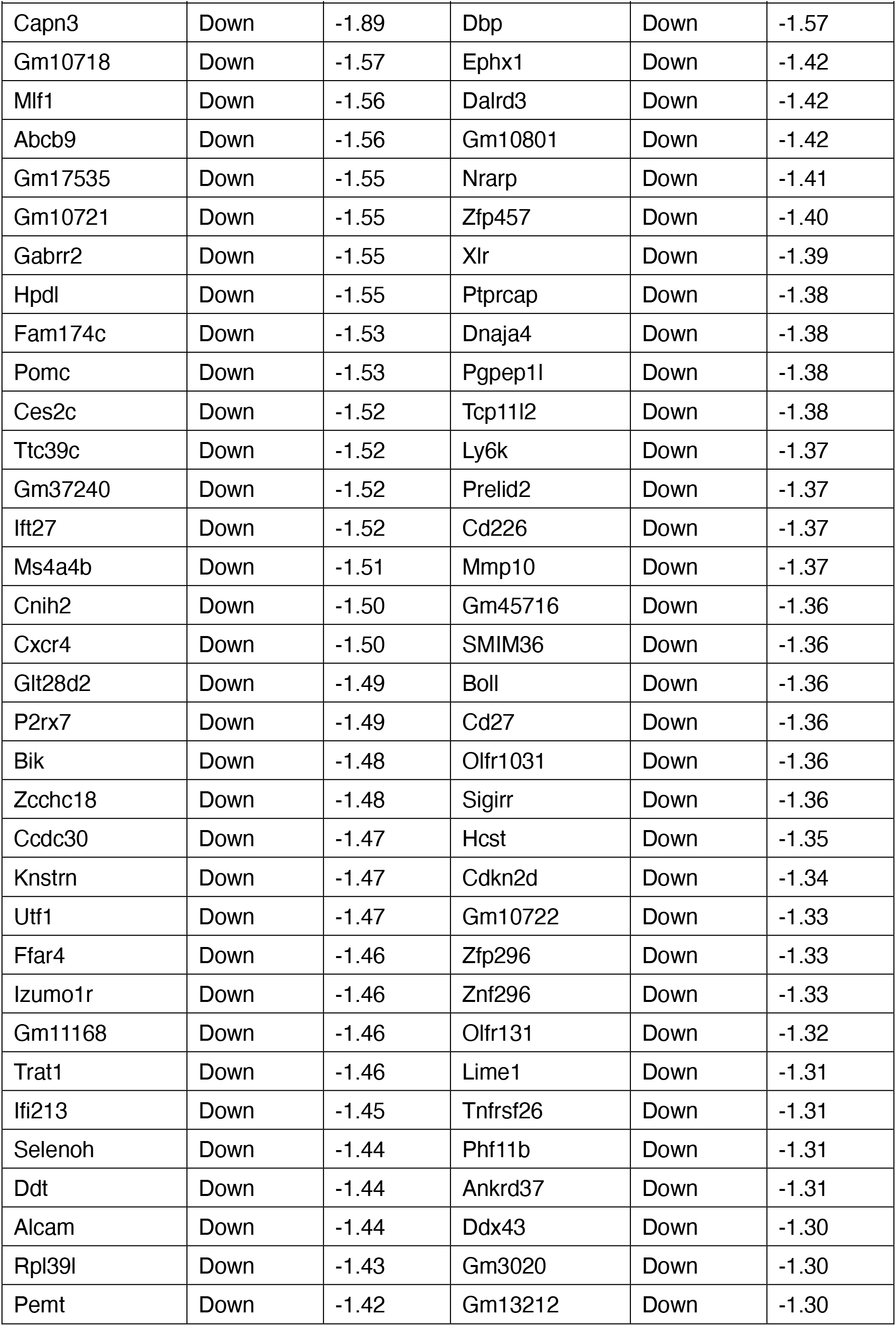
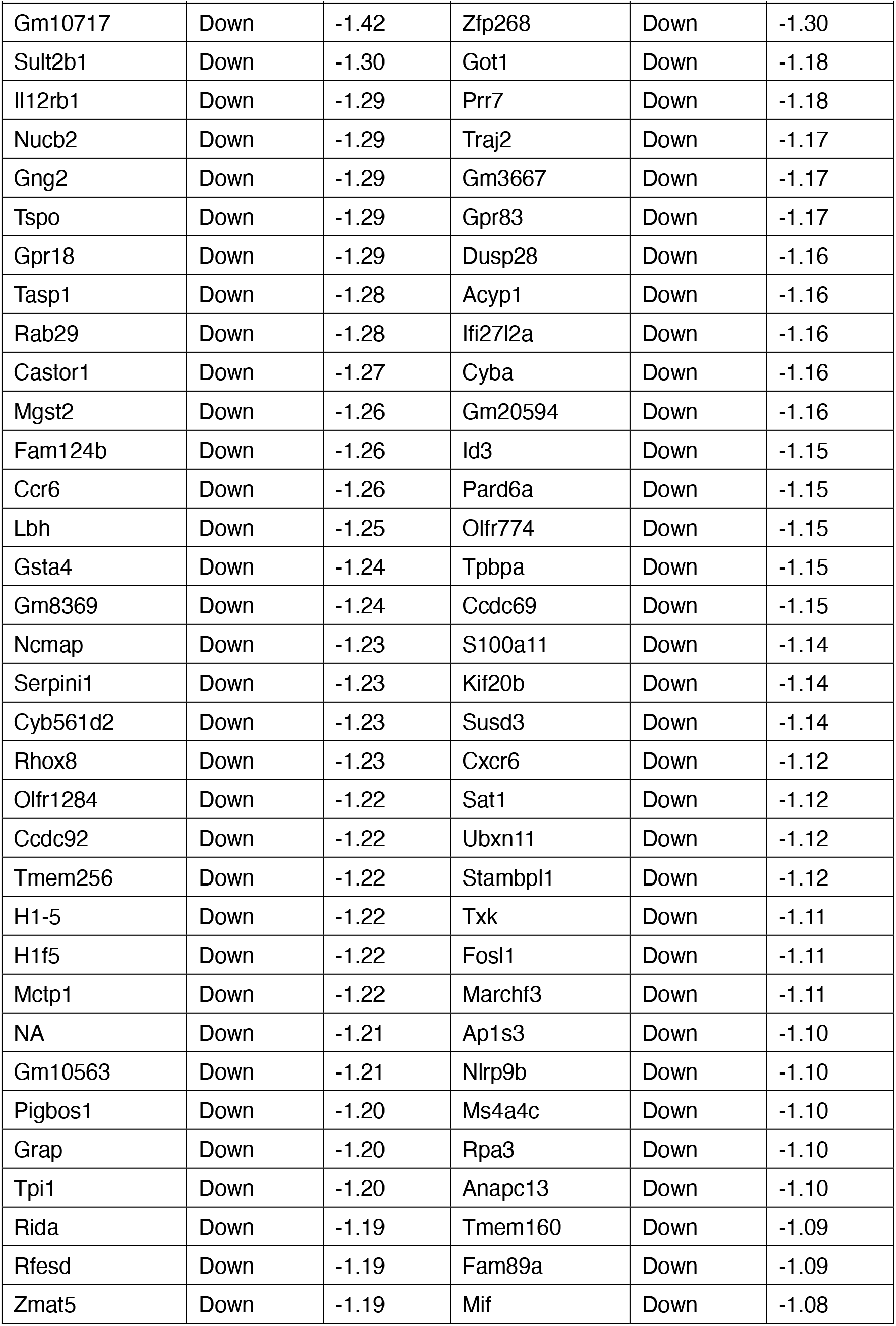
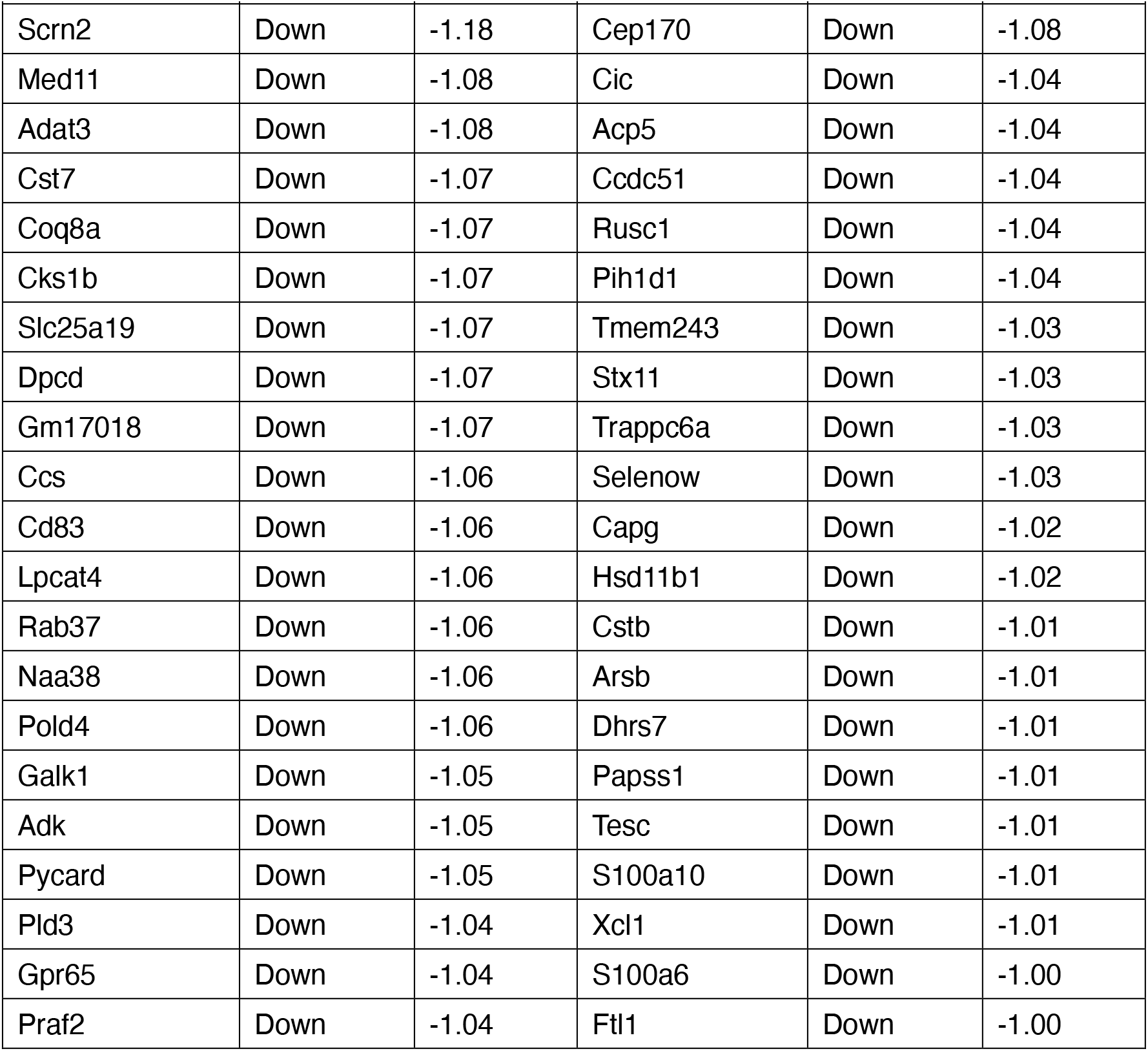
List of differentially expressed genes between wild-type and *IkE5*-deficient Treg cells.

